# Mice-side bias: Deliberative decision making in a model of rule revision reveals ‘myside’ confirmation bias-like cognitive processes in mice

**DOI:** 10.1101/2024.12.03.626648

**Authors:** Christopher Stevens, Cathy Lacroix, Mathilde Bouchet, Giovanni Marsicano, Aline Marighetto

## Abstract

Confirmation or ‘myside’ bias—over-valuation of novel information which confirms previously internalized cognitive content (prior beliefs, rules of conduct, etc.) and corresponding under-valuation of disconfirming novel information—constitutes a serious obstacle to adaptive revision of our beliefs, especially in ambiguous or complex epistemic environments. Indeed, myside bias has become a particularly pernicious fact of societal cohesion, contributing to the propagation of fake news, to social polarization, and even to the replication crisis in experimental science. By contrast, relatively little is understood about either its neurocognitive underpinnings or its evolution, one reason for this being that the potential presence of myside bias-like tendencies in non-human animals has never been directly tested. Hence, in order to advance research in both of these directions, we designed a novel mouse model of everyday-like rule revision such that the dynamic model environment would be sufficient to call out myside bias-like behaviors providing that mice did indeed possess the particular kind of competing neurocognitive processes necessary for it to manifest. Here, we both validate that model and provide the first behavioral descriptions of myside confirmation bias-like deliberative profiles in a non-human animal. Notably, we observe that this bias does not manifest in a merely unreflective or heuristic/‘system 1’ manner but rather also emerges through and indeed increases deliberative behaviors, especially in contexts of low representational resolution. Several other parallels with findings from human studies of myside bias are also detailed in the discussion.

## Introduction

In everyday language, when we say someone is ‘biased’ we are claiming that, as they explore and evaluate their environment, the mental image of it they construct is not a faithful representation but rather a simplified, skewed interpretation that fits a little too conveniently into their prior beliefs. This type of bias has classically been referred to as ‘confirmation bias’ and, as the authoritative review on the subject suggests, it can be observed everywhere, in many forms (Nickerson, 1998). Focusing more on the dimension whereby environment is interpreted to make it fit a given individual’s (e.g. ‘my’) subjective prior beliefs, this most commonly identifiable form of bias has more recently come to be known as ‘myside bias’ (Mercier & Sperber, 2017; Stanovich, 2021; Stanovich et al., 2013; Stanovich & West, 2007). Turning to its poorly understood underlying neurocognitive nature (Kaplan et al., 2016), a first question to ask is how such bias becomes instantiated in an agent in the first place.

A basic reinforcement learning (RL) way of thinking about this is that, as an agent/organism interacts with its environment, certain organism-environment interactions will be rewarded/positively reinforced, others will be value-neutral, while still others may be punished/negatively reinforced (Sutton & Barto, 2014). The expanding set of such reinforcement outcomes gives rise in turn to global associative conceptual systems representing what is useful, what is adaptive for a given organism to value, positively and negatively, in the environment it inhabits and must navigate. If the same conceptual value system is consistently reinforced, then, thanks to neurocognitive plasticity, it will become internalized by the organism, instantiated in its nervous system (Hebb, 1949; Mercado III, 2015). Thereafter, the organism will tend to prefer organism-environment interactions which are consonant with this internalized system of values and avoid or devalue those which are not (Festinger, 1957; Nickerson, 1998). Importantly, this preference and avoidance is not a mere matter of coldly calculating predicted reward values but furthermore implies emotional/affective components which also guide behavior, particularly in situations of cognitive uncertainty (Damasio, 1994; Bechara & Damasio, 2005; Damasio, 2012; Panksepp, 1992, 1998). Ultimately, this cognitive and affective guiding of behavior imposes certain constraints. It ‘biases’ the organism. Stated plainly, once we have internalized such a system, “it is neurally instantiated in our brains and we are not free to think just anything” (Lakoff & Johnson, 1999). The particular lack of cognitive ‘freedom’ associated with myside bias has emerged as an undermining factor of the information age, blamed for contributing to the propagation of fake news (Lazer et al., 2018), to the polarization of society into mutually hostile camps (Del Vicario et al., 2017; Stanovich, 2021), and even to science specific dilemmas such as the replication crisis (Baker, 2016; Nuzzo, 2015).

What are the cognitive processes through which myside bias produces such effects? First, given a particular internalized/instantiated conceptual system, relevant information from the environment becomes either *confirmatory* or *disconfirmatory* with respect to it. Second, classical confirmation bias theory advances that confirmatory information will be *over*valued by the organism and *dis*confirmatory information *under*valued, e.g. explained away (Nickerson, 1998). This characterization explains why confirmation bias penetrates almost every aspect of life in which new information from the environment has bearing on prior beliefs (Nickerson, 1998). Indeed, the occurrence of myside bias may actually be augmented by quantitative increases in available information for the reason that such increase provides more ‘raw material’ for biased sampling and evaluation (Stanovich, 2021). Finally, a widely misunderstood but crucial point about myside bias is that its strength in a given individual does *not* correlate with standard measures of general intelligence (Stanovich et al., 2013; Stanovich & West, 2007). Indeed, as far back as Francis Bacon it has been noted that scientists and philosophers are frequent exemplars of biased reasoning (Bacon, 1620; Nickerson, 1998; “The virus of myside bias is spreading among cognitive elites - Interview with Keith E. Stanovich,” 2020).

A core reason for our relatively poor neurocognitive understanding of myside bias (Kaplan et al., 2016) is an absence of animal models designed to search for and investigate analogous cognitive processes in non-human species. Taking this as our aim, we designed a novel mouse model of rule-revision such that, if the murine organism does indeed comprise the necessary neurocognitive processes, then the protocol will be sufficient to elicit myside bias-like behavior in mice. Using the 8-arm radial maze throughout, the model consists of two steps: 1) An initial rule-training phase to ‘bias’ cognition (modeling initial instantiation of a conceptual system): 2) A subsequent rule-revision phase designed to generate both confirmatory and disconfirmatory information with respect to the initially internalized rule (modeling environments liable to elicit myside confirmation bias). (Note that ‘rule-revision’ is to be understood as functionally synonymous with ‘belief revision’, insofar as both beliefs and rules constitute policies for determining action decisions in given states (Peirce, 1878), i.e. what in RL are called “state-action policies” (Sutton & Barto, 2014).)

In the first phase, mice underwent an intense learning schedule to robustly internalize a putatively striatal (McDonald & Hong, 2004) tactile stimulus-response (S-R) behavioral rule (R1). Each trial of this phase consisted of a choice between two surfaces, one always predictive of reward location (S1), the other reward absence (S0). In the second phase, which we call everyday-like rule-revision (EdRR), R1 trained mice were introduced into a highly novel radial maze environment. Here, in order to successfully perform a hippocampus-dependent, everyday-like memory (EdM) task (Al Abed et al., 2016; Stevens et al., 2023) wherein reward location is determined not by a tactile state-action policy but rather a spatial alternation-based one (Dember & Richman, 1989), mice had to simultaneously inhibit and revise R1. Crucially, by adding the S1 and S0 surfaces to the classical EdM experimental design, every EdM trial outcome also constituted either a confirmation or a disconfirmation of the R1 rule. Furthermore, the EdM protocol also entails trials of varying mnemonic difficulty, resulting from how many times the memory episode of a given trial *n-1* has been representationally weakened via repeated organizational ‘active forgetting’ (Bekinschtein et al., 2018; Hulbert et al., 2016) prior to being required for accurate response on the subsequent spatially relevant trial *n* (figure 1C; for full details see Stevens et al., 2023). Moreover, this feature of the EdM protocol is the reason we chose it as the basis for our model of myside bias: given that, in human studies, myside bias increases as a function of the (subjective) cognitive complexity of a given epistemic situation (Stanovich, 2021), an animal model of the phenomenon should similarly give rise to bias that increases as a function of cognitive uncertainty. Indeed, the aim behind our design was to introduce a model of real-world, often ambiguous situations characterized by belief intrusion and cognitive interference, in contrast to classical ‘pure’ reversal learning protocols neither suitable for generating sequences of confirmations and disconfirmations nor sensitive to varying degrees of cognitive uncertainty. Finally, we must underline that this experimental design rests upon a profoundly consequential twofold cognitive assumption. First, that the observable behaviors by which we identify myside bias in humans result both from initial neural instantiation of conceptual systems *and* from particularities of the (epistemic) environments our species has created for itself and must navigate. Second, that there is therefore no *a priori* empirical reason to presume that the underlying neurocognitive components necessary for myside bias to manifest under particular environmental conditions are a phylogenetic novelty of humans (as has recently been claimed by Mercier & Sperber, 2017).

**Figure 1.**
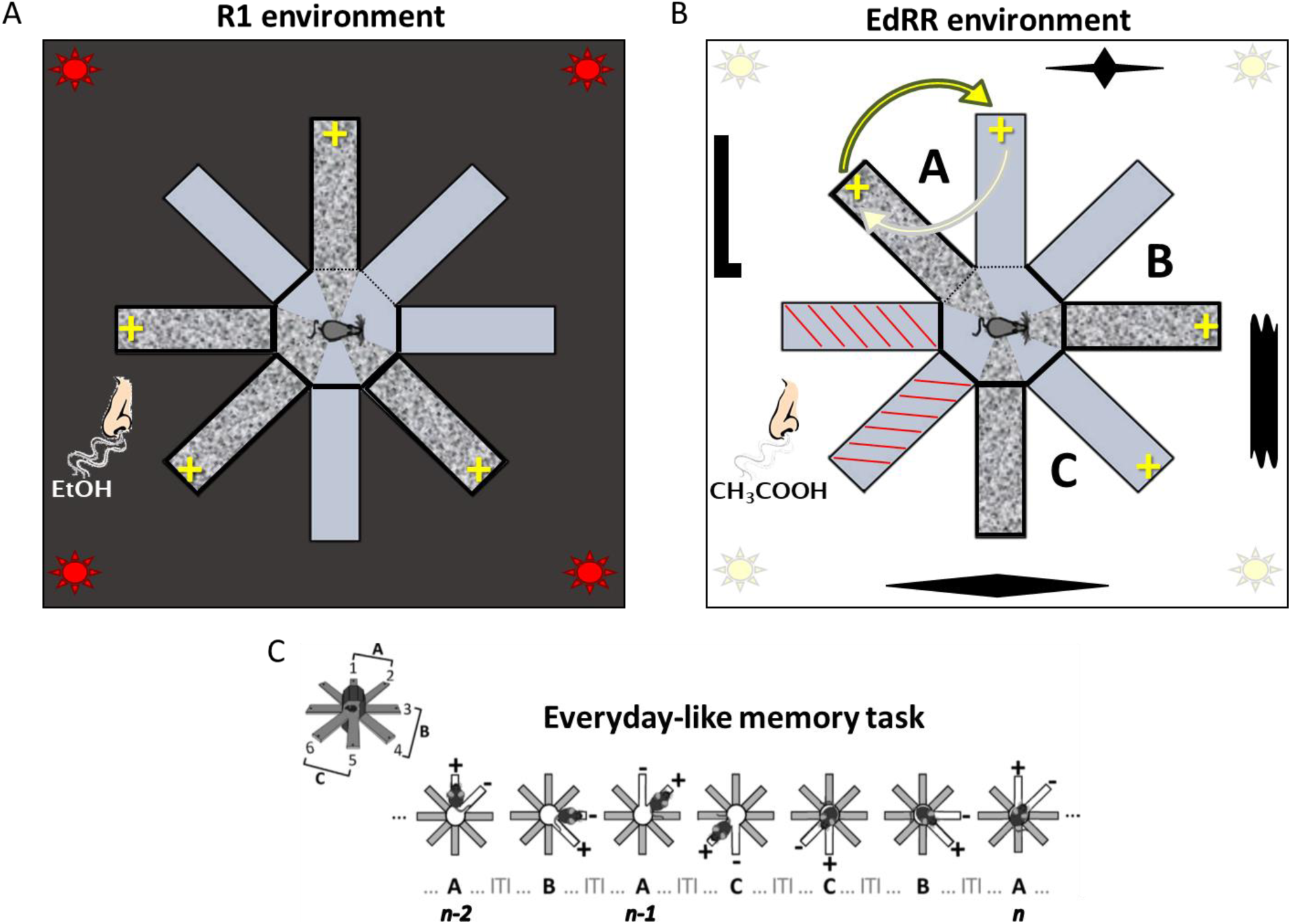
Behavioral environments and tasks. Graphical representation of the radial maze environments composing the full everyday-like rule revision protocol. **(A)** The R1 tactile discrimination phase is conducted in darkness, removing all visual spatial bearings, with the objective of focusing attention on the tactile dimension. The surface of the radial maze is divided in two according to surface: a smooth surface, represented in solid gray, and an irregular surface, represented in dappled gray. Each animal learns that one of the two surfaces is always predictive of food reward location (yellow ‘+’ symbols). This surface is referred to as S1. Whether S1 is the smooth or the irregular surface is counter-balanced among the animals. All 8 arms are used in this phase. Each trial consists of a choice between two contiguous arms, one S1, one S0. A 70% ethanol mix is also used to give this R1 environment a distinct olfactory dimension. **(B)** Once animals have completed R1 training (either by reaching 75% criterion level averaged across two sessions or by completing a fixed number of R1 sessions, depending on experimental conditions; see main text) they are moved to the EdRR radial maze environment, which contains several novel differences with respect to the R1 environment: radial maze situated in a separate room; lights on, hence visibility of spatial landmarks (represented by the black shapes around the radial maze); 1% acetic acid mix used to distinguish environment at the olfactory level. The EdRR task, similar to the classical EdM task, uses 6 arms of the radial maze, arranged into three pairs, A, B, and C. However, EdRR is distinct from EdM by the presence in each pair of one S1 and one S0 arm. The cognitive consequence of this is that every EdM trial (spatially alternate relative to previous choice on present pair; see below) simultaneously constitutes an instance of an R1 trial (choose S1 rather than S0). This allows us to qualify each EdM error made during performance of EdRR as either an R1/S1 or a nonR1/S0 type error, providing the basis for the quantification and analysis of R1 bias during everyday-like revision of R1. **(C)** Illustrative sample sequence of classical EdM task. On trial *n-2*, the animal chose the left arm of pair A and was rewarded. Trial *n-1* on pair A then constitutes a level 1 trial, as there has been 1 interposed trial on pair B between *n-2* and *n-1*. The animal correctly chooses the right arm on *n-1*. When pair A is next presented, it constitutes a level 3 trial, as there have been 3 interposed trials (pairs C, C, and B) between *n-1* and *n*. To choose correctly on trial *n*, the animal must focus on recalling which arm it chose on trial *n-1*, inhibiting interference from both the interposed trials on other pairs and from the memory of trial *n-2*.

Mice performing EdRR reflected the above-mentioned myside bias behaviors in several ways. The majority of errors committed during EdRR occurred on/were biased towards the S1 surface but, when compared to control animal performances, no prominent impact on overall number of EdM errors (representing ‘general intelligence’) was observed. Initial exploration of the novel environment was also not diminished, indicating that the primary cognitive consequence of prior learning was not reduction in sensitivity to novelty *per se* but rather an imbalance in subsequent *evaluation* of novel environmental information. The probability of choosing S1 rose significantly as a function of increasing EdM trial difficulty, indicating that increased active forgetting-induced cortico-hippocampal cognitive uncertainty entailed higher likelihood of reverting to the more habitual, putatively striatal R1 response strategy. Analyses of fine-grained deliberative behavioral measures (Stevens et al., 2023) also revealed that bias did not manifest primarily as a fast ‘system 1’-like heuristic (Kahneman, 2011) but exerted influence *through deliberation*, which in fact it increased. Based on our results, we suggest that R1 bias and its inhibition are partially *independent* of the cortico-hippocampal functions necessary for EdM performance, again commensurate with the lack of correlation between myside bias and general intelligence documented in humans. To conclude, we briefly outline a multiple memory systems interpretation of our results (McDonald & Hong, 2004; McDonald & White, 1993; White & McDonald, 2002), advancing that tasks like EdM/EdRR are uniquely suitable for investigating the balance between the respective contributions of multiple systems and, ultimately, how this balance is impacted by environment, cognitive dysfunction, ageing, etc.

## Materials & Methods

### Animals

Young (8 to 12 weeks) C57BL/6J male mice were obtained from Charles River and collectively housed in a standardized animal room (23 °C; lights on 7 AM to 7 PM; 4 to 5 mice per cage). All animals were moved to individual cages 2 weeks before the beginning of experiments.

### Food restriction

5 days prior to the first day of training, all animals were placed under a progressive food restriction schedule in order to gradually bring them to 85% to 90% of their baseline free feeding weight. Individual animal weight and welfare was monitored daily throughout the duration of the experimentation. All experiments were conducted in accordance with European Directive 2010-63- EU and with approval from the Bordeaux University Animal Care and Use Committee CCEA50. All efforts were made to minimize suffering and reduce the number of animals used.

### Radial maze

8-armed fully automated radial mazes (Imetronic), whose surface is raised ∼100cm off ground level, were used. Access to each arm is from a central platform by means of automated vertically retracting doors. In-task, when all doors are closed, the experimental animal is contained within the central platform, a regular octagon of area ∼485cm² and edge 10cm (i.e. the width of each arm and door). At the distal end of each 50cm length arm is an automated pellet distributor for dispensing food reward. The distributor is set into a slight indent in order to hide its state (i.e. baited/not baited) from the animal. For this study, we produced removable polymer panels which could be placed so as to cover the entire area of the radial maze. The panels, painted the same color as the maze, were of two distinct tactile finishes, smooth surfaced panels (similar to the usual surface of the radial maze) and irregular surfaced panels (the finish of which was a uniform but irregular beveled pattern of < 2mm maximum relief). This allowed us to present the radial maze according to various tactile configurations: entirely smooth, entirely irregular, or some combination of smooth and irregular. Animal movements in the radial maze were detected via video camera and motion detection software (GenCam) using either visible or infra-red light, depending on the experimental conditions (see below). The motion detection software communicates with a second piece of software, POLYRadial (Imetronic), through which pre-programmed sequences of automated radial maze actions are triggered. This program was used for the design and execution of behavioral exercises (sequences of door openings, location of food reward, conditions for opening and closing of doors, etc.). Hence, the exercises were contingent upon a combination of both the detected movements of the animals plus automated timed sequences.

### Behavior

#### Habituation

Prior to the first day of tactile discrimination learning, all animals were habituated to the context and functioning of the radial maze apparatus. Food restriction, as described above, began 3 days before habituation (i.e. 5 days before training). At the beginning of each habituation session, the animal was placed by the experimenter in the central platform of the radial maze, all 8 doors of which were closed. Once removed to the control room, the experimenter launched the habituation program via the POLYRadial software. The program began by an interval of 10 seconds during which the animal could explore the central platform. Following this, all 8 doors opened simultaneously, presenting the animal with the opportunity to freely explore the entire surface of the maze. As the animal explored, once it had advanced to the most distal section of a given arm (location of the food reward distributor) and returned to the central platform, the door of that arm automatically closed behind it, thus preventing further access in the current session. Once the animal had fully explored all 8-arms, it thus found itself again contained within the central platform. Further habituation sessions could then be launched by the experimenter if necessary. It was considered that when at least 5 out of 8 available food rewards had been recovered and consumed in a single session, the animal was fully habituated to the relevant functionalities of the apparatus. All animals reached this habituation criterion within an average of 5 sessions. Since the tactile discrimination phase was to be conducted in the dark, rendering it potentially difficult for animals to perceive and learn that doors always open in contiguous pairs, animals also underwent a second habituation session 24 hours after the first. Here, pairs of doors opened simultaneously, creating a choice to explore one of two neighboring arms, as would be the case in the subsequent tactile discrimination phase, but both arms were rewarded. Crucially however, during both phases of habituation, the surface of the radial maze was *entirely* covered in the surface type (smooth or irregular) already assigned to be a given animal’s rewarded surface during the subsequent tactile discrimination training. In this way, experimental animals had zero experience of being rewarded on what would be their assigned *un*rewarded surface during tactile discrimination training.

#### Tactile discrimination

In a first phase, mice were trained in the radial maze to discriminate between two surface types, smooth and irregular, in a stimulus-response (S-R) manner. For each animal, only one of the surface types was predictive of reward location (S1) and the other predictive of reward absence (S0). Ethanol (70%) was also used to add a specific odor-based dimension to the R1 environment. This initial phase was conducted in darkness to oblige reliance on non-visual (notably tactile) sensory inputs. Each trial consisted of choosing which of two neighboring arms (one S1, one S0) to visit. Mice were considered to have fully internalized the S1-reward association rule (R1) when they had reached a mean performance of 75% correct responses averaged across two sessions. Control animals were rewarded on every trial during this phase, regardless of which surface they chose. Once experimental animals reached criterion level performance in the R1 environment, they were moved to a new radial maze in a new room, in presence of a new odor (acetic acid, 1%), with new lighting conditions (lights turned on) rendering intra- and extra-maze cues visible. The rationale for these contextual changes was to emphatically announce, via multiple sensory systems, that a *novel* environment, distinct from the R1 environment, had been entered (see *Rule Revision* below).

#### Everyday-like memory

As previously described (Al Abed et al., 2016; Stevens et al., 2023), the aim of the EdM radial maze protocol is to model daily life situations in which sets of contextually related memory episodes compete for attention and must be organized via ‘active forgetting’ (i.e. ‘forget about that for a moment’) to enable accurate episode retrieval and subsequent responding in given spatiotemporal task contexts.

Responding to the EdM challenge (figure 1B), animals must simultaneously: 1) Mnemonically encode spatiotemporally distinct episodes regarding the action decisions they are making and outcomes they are experiencing on three distinct task contexts (arm pairs *A*, *B*, and *C*). 2) Continuously organize all of these mnemonic episodes in order to choose the correct action on subsequent trials (according to a simple rule of ‘always choose the action *not* chosen during the previous trial on a given pair’; figure 1C).

In short, the task implies mnemonically encoding and storing information relative to which arm was visited on any given arm pair presentation *n-1* until the next trial *n* consisting of a presentation of the same arm pair (figure 1C). The task relies on and reinforces the spontaneous mouse behavior of spatial alternation, implying a spatial ‘win-shift’ strategy (McDonald & White, 1993) as opposed to the tactile ‘win-stay’ strategy required for R1 performance. The number of interposed trials (between 0 and 4) on either of the other two task contexts constitutes the variable retention component of the task (5 levels of difficulty, 0 to 4). However, retention in this task is highly dynamic and, for every EdM trial, a significant organizational mnemonic component is present in both the necessity to inhibit (‘actively forget’) spatially irrelevant content pertaining to the *n-1* trials on the *other* two pairs as well as the need to inhibit spatially relevant but temporally (or episodically) *ir*relevant interference on any given arm pair presentation *n*. This is to ensure that the previous *n-1* choice and not the previous-previous *n-2* choice on the present pair is acted upon (figure 1C). Both these varieties of trial *n* context-irrelevant memory episodes constitute interfering and competing cognitive noise in the decision making process. Level 0 difficulty trials correspond to two immediately consecutive presentations of the same arm pair, with no interposed trials on the other pairs. Level 0 thus most closely resembles a classical T- or Y-maze spontaneous alternation trial (figure 1C; the second presentation of pair *C* is a level 0 trial; the second presentation of pair *A*, a level 1 trial; the third presentation of pair *A*, a level 3 trial, etc.). However, in contrast to classical T- or Y-maze contexts, even on level 0 trials, there may be cognitive interference from episodes irrelevant to the presented arm pair.

EdM performance can be analyzed session by session, block by block (each block = 3 sessions and is comprised of 12 trials of each trial difficulty level, facilitating more robust statistical analysis), or averaged across all sessions/blocks according to trial difficulty level. For full details see (Stevens et al., 2023).

#### Rule revision

The primary specificity of the everyday-like rule-revision (EdRR) task consists in the presence, on each arm pair and during each EdM trial, of one S1-covered and one S0-covered arm, counter-balanced left and right between pairs. This tactile modification of the EdM environment entails that, as an animal performs the EdRR task, a sequence of confirmations and disconfirmations of R1 is generated: whenever a S1 arm is rewarded or a S0 arm is not rewarded, R1 is *confirmed*. Conversely, whenever a S0 arm is rewarded or a S1 arm is not rewarded, R1 is *disconfirmed*. It is on this basis that we present this rule revision model as a potential animal model of myside bias.

In brief, as R1 trained mice perform the EdRR task, they not only must retain and organize their previous actions as per the classical active forgetting-dependent EdM task, but additionally they must actively inhibit the R1 rule in order to ensure that they respond to each trial as an instance of EdM and not as an instance of R1 (obeyance of which would lead to them repeatedly choosing S1 instead of spatially alternating). This addition of S1 and S0 to the EdM environment enabled us to qualify each choice and error an animal made. We could then calculate both the total number of S1 choices and the ratio of S1 errors to total errors, providing us with two strong measures for assessing the magnitude and evolution of R1 interference/bias in EdM behavior.

Note that, in contrast to the classical EdM task where both arms are baited on the initial trial in each session, in EdRR only the S1 arm is rewarded on these three initial trials, following which reward location spatially alternates as described above. In other words, the optimal action policy for the EdRR task can be formalized as follows: 1. On the initial trial of each pair in a given session, choose the S1 arm. 2. On all subsequent pair presentations, spatially alternate with respect to previous choice.

### Analysis

All raw data extraction, analysis, statistical comparison, and graphical representation was generated using custom codes written in Python (Van Rossum & Drake, 2009) using the pandas (Reback et al., 2020), numpy, pingouin (Vallat, 2018), bioinfokit (Bedre, 2021), matplotlib (Hunter, 2007), and seaborn (Waskom, 2021) libraries. All code is open source and available at https://github.com/metaphysiology.

### Glossary of deliberative behaviors

#### Decision latency

Time elapsed (in milliseconds) between the instant when a trial begins (doors of the current trial pair open) and the instant when the threshold from the central platform into the surface-arm of the animal’s definitive choice is first crossed.

#### Run time

Time taken (in milliseconds) for animals to travel the distance from the threshold of the definitively chosen arm to the reward-distributor containing distal zone.

#### Choice revision

Number of times (units per trial) an animal crossed the threshold into one arm of a pair only to revise their choice by exiting it again prior to entering its most distal zone (which would trigger the door to the unchosen arm to close). At this point animals could either choose to explore the other arm of the pair or (more rarely) re-enter the same arm. As long as the distal zone of either arm had not been entered, this process could technically continue indefinitely. Each choice revision was quantified as a ‘KOOK’ unit, capturing the fact that some choice revisions were ultimately error-inducing, ‘KO’, while others were rectifying, ‘OK’.

For more detailed discussion of all three deliberative parameters, see (Stevens et al., 2023).

## Results

Young C57Bl6/J mice completed 12 sessions/4 blocks (see Materials & Methods) of the everyday-like rule-revision task (EdRR). This number of sessions allowed us to characterize not only the initial and overall cognitive impact from the internalized R1 rule but also its evolution over time/repeated everyday-like memory (EdM) training.

Across three iterations of the experiment, animals were divided into two groups: animals who were trained in the first environment to acquire the R1 tactile discrimination rule up to criterion level (R1 population, n = 40), and; animals who were rewarded on every trial in the first environment, regardless of which surface they chose (controls, n = 19). Reward location in the classical EdM task is determined by trial-by-trial spatial alternation between the arms of each given pair (total 3 pairs; *A*, *B*, and *C*, see figure 1B). During EdRR, each trial of the EdM task also constitutes an instance of the prior R1 choice between visiting an S1 or an S0 arm. In order to contextually mark the transition between the R1 and EdRR tasks, we designed the environments so that the second would be as novel as possible with respect to the first: new room and radial maze, change of visibility conditions from darkness to lights on, consequent presence of visual spatial landmarks, plus change in ambient odor (figure 1A-B).

### 1. Prior learning impacts quality more than quantity of EdM errors

#### 1.1 Weak specific impact of prior learning on everyday-like memory performance

R1 trained and control animals performed comparable levels of correct responses per session and displayed comparable learning curves over the 12 sessions/4 blocks of training, with significantly more errors observed in the R1 population only in block 1 (sessions 1-3) (figure 2A; one-way ANOVA with pairwise Tukey post hoc, block 1: *F*(1, 175) = 4.9, *p* = 0.028). Moreover, individual R1 population performances in the EdRR task did not correlate with either averaged or weighted averaged (applying discount factor to earlier R1 sessions) R1 phase performances (supplementary figure S.1A, *R*² = 0.002; supplementary figure S.1B, *R*² = 0.00).

**Figure 2.**
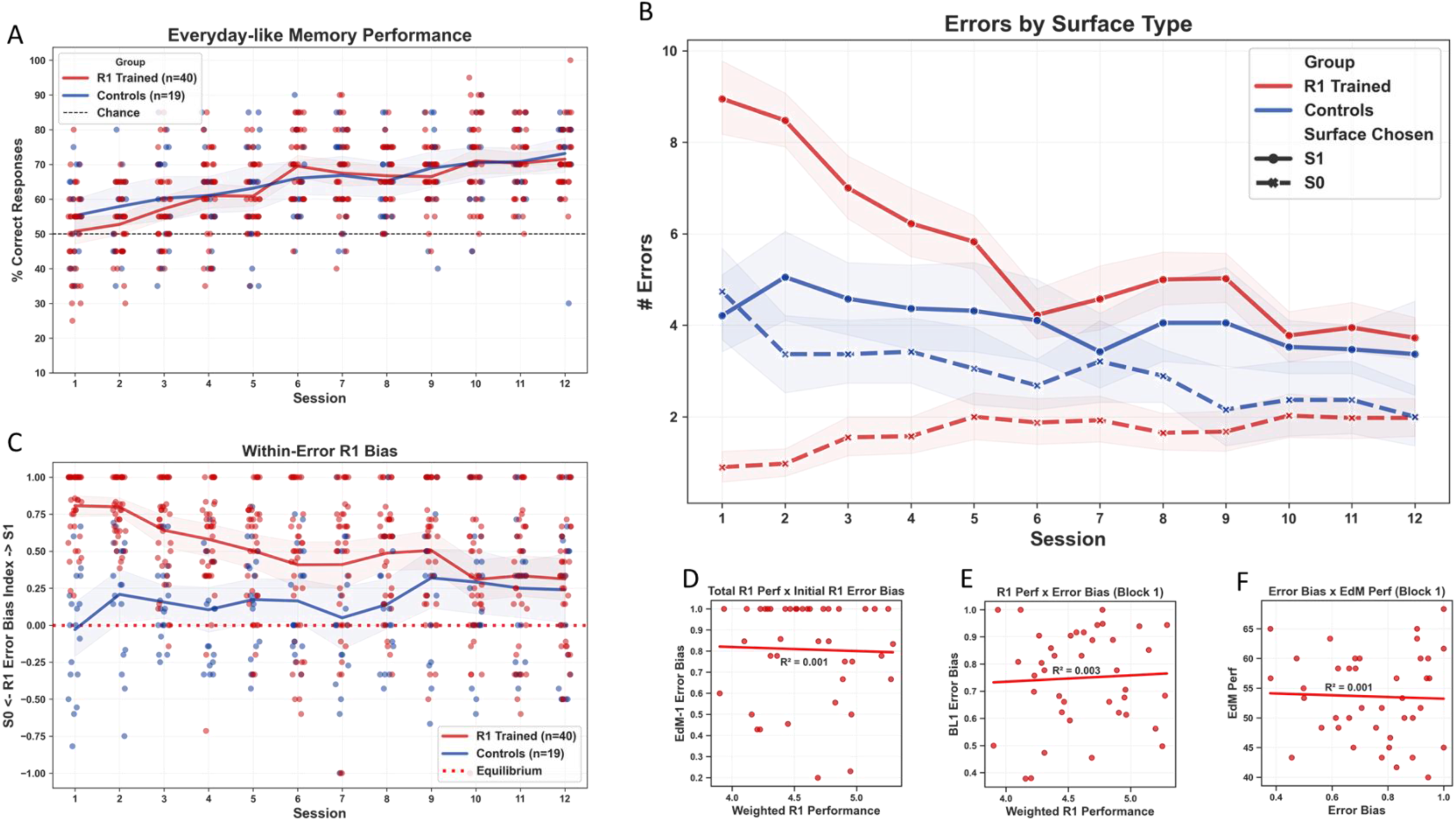
Impact on everyday-like memory of a previously learned rule especially visible in quality of errors committed. R1 animals are represented in red (n=40), controls in blue (n=19). All error bands represent 95% confidence intervals, vertical spaces between bands provide visual indication of statistical significance; detailed statistical analysis in main text. Curves represent population means, dots are individual values. **(A)** Everyday-like memory performance calculated from 20 per session trials. R1 trained and control populations both began with very low, barely above chance level overall performances which improved gradually with repeated training and at a comparable rate. Controls performed slightly but significantly better than R1 trained during sessions 1-3/block 1 only. **(B)** Although both populations made similar overall numbers of EdM errors, during EdRR 1 + 2 especially, ∼90% of R1 trained errors were biased towards S1. This proportion decreased with repeated EdRR training, primarily as a function of decreasing numbers of S1 errors. Even in final EdRR sessions, however, significantly more errors occurred on S1 compared to S0. Control animals also displayed a slight emergent bias towards S1, significant when averaged across all sessions but not in any given session (analyzed further below). **(C)** We then calculated a within-error R1 bias index per animal, per session by taking the relative difference between S1 and S0 errors (S1 errors – S0 errors / Total errors) and tracking its evolution. No significant decrease was observed between EdRR 1 and EdRR 2 in R1 trained animals, after which the within-error R1 bias began to gradually decrease, though even after 12 sessions of EdRR errors were still biased towards S1. Controls also developed a slight S1 error bias over repeated EdRR training, which achieved statistical significance in the final 4 sessions. **(D)** + **(E)** Mirroring human studies showing no correlation between strength of myside bias and overall cognitive ability, linear regression analyses revealed no correlation between strength of individual performance levels in the R1 training phase and levels of within-error R1 bias in either session 1 or block 1 of EdRR. **(F)** Linear regression analysis also found no correlation between level of within-error R1 bias and overall EdM performance across block 1, indicating that how biased an animal was in the errors it committed was no predictor of how many errors overall it would make, and vice versa, once again very closely mirroring findings in human studies of myside bias.

#### 1.2 Strong impact of prior learning on within-error R1 bias

Beginning with the first session (EdRR 1), though similar in terms of mean total number of errors (EdRR 1: R1 population, mean # errors 9.85; controls, mean # errors 8.95), the quality of these errors was vastly different between the two groups. During EdRR 1, the mean ratio of errors on S1 to S0 arms was roughly 9:1 in the R1 population, whereas in controls it approached 1:1 (4.2:4.7) (figure 2B; mixed ANOVA design, interaction between ‘Group’ within ‘Surface’: *F*(1, 57) = 47.4, *p* < 0.0001). Even during EdRR 2, conducted 24 hours after EdRR 1, this mean error ratio decreased only very slightly in the R1 population, to around 8.5:1. We also analyzed and represented this error bias and its evolution over repeated EdRR training as a relative difference ((S1 errors – S0 errors) / Total errors), calculating per animal and per session what proportion of its total errors were biased towards the R1 rule. This revealed that in many R1 trained animals, especially in the early EdRR sessions where the greatest total number of errors were made, 100% were committed on S1 arms (figure 2C). Overall, the R1 population made a significantly higher proportion of S1 errors than controls (figure 2C; one-way ANOVA with pairwise Tukey HSD post hoc: *F*(1, 705) = 96, *p* = 0.001). No significant decrease in this within-error R1 bias was observed between EdRR 1 and EdRR 2. We did also observe an emergent trend for controls to commit a comparatively higher proportion of errors on S1 arms than on S0 arms, though this was significant only when averaged across multiple sessions (repeated measures ANOVA across all sessions: *F*(1, 18) = 20.5, *p* < 0.0003) and will be further analyzed below.

This first set of results demonstrates that the strongest and most persistent impact of prior R1 training was manifest not in EdM performance itself but rather in the type or *quality* of the errors committed. However, linear regression analyses revealed no correlation between individual strength of within-error R1 bias in EdRR 1 and prior final R1 phase performances (supplementary figure S.1C, *R*² = 0.018). Nor was a correlation found when comparing total weighted R1 performance (using a discount factor to give more weight to later compared to earlier R1 sessions) against within-error R1 bias during either EdRR 1 (figure 2D, *R*² = 0.001) or block 1 (i.e. EdRRs 1 – 3; figure 2E, *R*² = 0.01). Finally, individual within-error R1 bias index values also did not correlate with EdM performance in block 1 (figure 2F, *R*² = 0.001). Taken together, we interpret this absence of simple correlations as preliminary evidence that the behavioral output elicited by the EdRR task results from complex interactions between multiple cognitive systems, the relative ‘strengths’ of which are, to some meaningful degree, independent of each other.

### 2. As novelty decreases, bias increases

#### 2.1 Novelty triggers exploration

As just seen, the R1 population displayed marked within-error R1 bias in the EdRR task without, however, committing many more total errors than controls. At the most basic level, this is because they did not fully avoid S0 arms. By analyzing intra-session trial-by-trial behavior, we were furthermore able to observe that, in EdRR 1 especially, the primary driver of S0 arm exploration was not, as may have been expected, an accumulation of unrewarded visits to S1 arms triggering a subsequent tactile ‘lose-shift’ strategy. Rather, upon initial placement in the highly novel EdRR environment, the R1 population explored all available arms, S1 and S0, on average even more efficiently than controls (figure 3A). To quantify this behavior, we calculated an exploration index (EI) for each animal, lower values representing quicker exploration of all 6 available arms. EI was therefore calculated as a contingent function of the minimum number of trials required in a given EdRR session trial sequence for all 3 arm-pairs to be presented at least twice. In EdRR 1, for example, on trial number 7 an animal will have been presented with all three arm-pairs twice, providing it with sufficient occasion to have explored all arms once each. Hence, animals who explored all 6 arms by trial number 7 of EdRR 1 were attributed an optimal EI of 0, labelled ‘Optimal’ on the y-axis in figure 3A (optimally exploring animals in EdRR 1: R1 trained, 20/40, 50%; controls, 6/19, 31.6%). Animals who explored all 6 arms at least once by trial 8 were attributed an EI of 0.0625 (R1 trained, 7/40, 17.5%; controls, 3/19, 16%), and so on. Only 2/40 (5%) R1 trained animals failed to explore all 6 arms of the novel radial maze by the end of EdRR 1. These animals were attributed an EI of 1.1, labelled ‘Fail’ on the y-axis of figure 3A. Overall in EdRR 1, the R1 population had a median EI of 0.0625, i.e. one degree above optimal, whereas the control group had a median EI of 0.1875, though no statistically significant effect of ‘Group’ was found. We take these results as demonstrating either that the high initial novelty of the EdRR environment amplifies the exploratory drive or, we instead suggest, that it inhibits ongoing R1- driven active inhibition of the exploratory drive thereby *disinhibiting* it (see below). Importantly, both interpretations can explain why the impact of novelty on exploration was more prominent in the R1 population than in controls, since for R1 trained animals the ‘explore’ imperative translates operationally as ‘choose S0’ as opposed to ‘choose S1’, the latter corresponding to the S-R ‘exploit’ command they had been trained to act under during the R1 phase.

**Figure 3.**
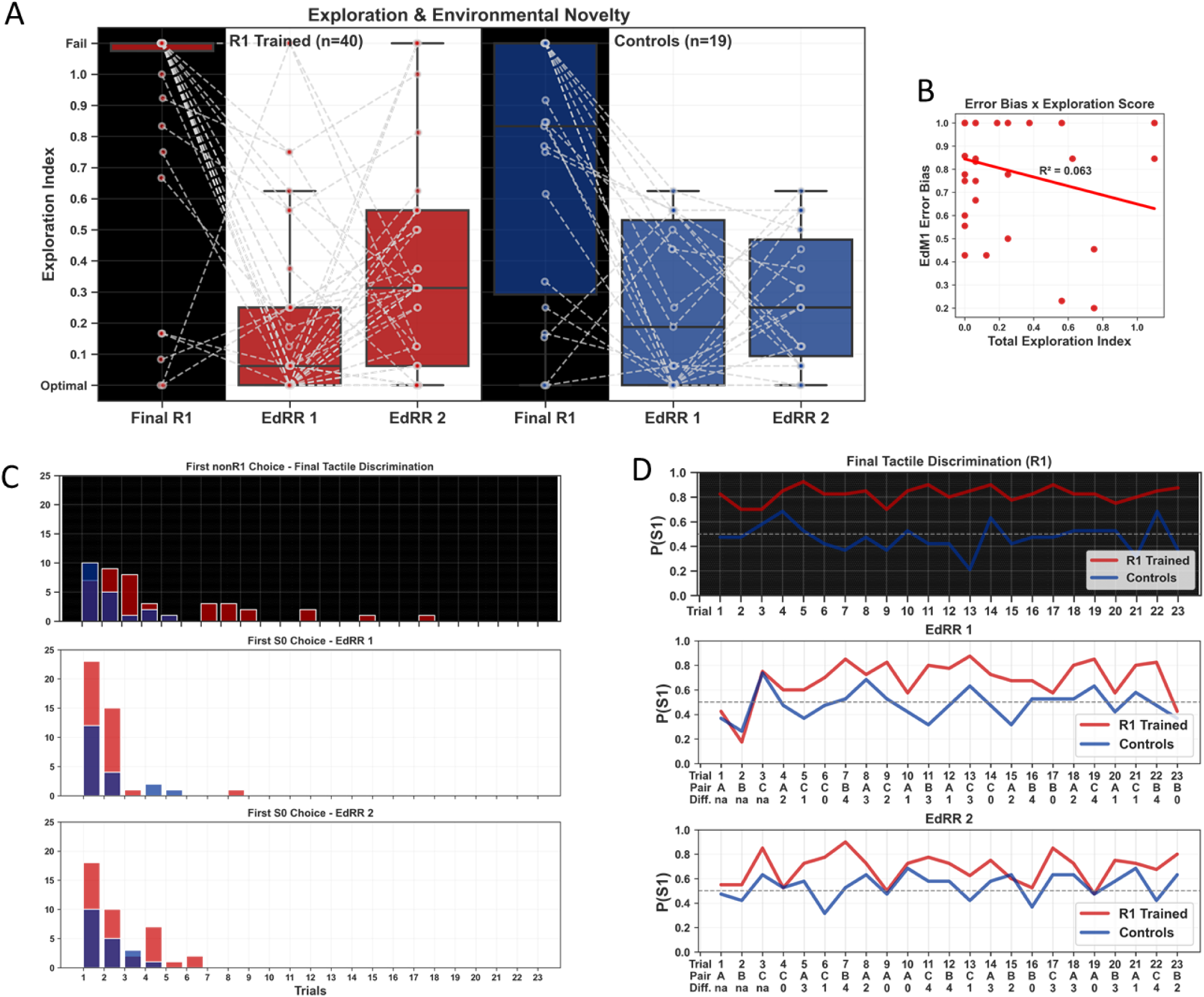
High initial environmental novelty-driven exploratory behavior decreases as novelty fades. Transfer from the R1 to EdRR environment constitutes high multi-sensorial novelty (see figure 1). Here we assess the impact of this novelty on behavior. R1 trained animals (n=40) represented in red, controls (n=19) represented in blue. **(A)** Exploration index (EI) per animal per session, calculated from number of trials (as a function of total number of trials) within which animals explored each available arm at least once. Low EI values indicate all available arms were explored within a low number of trials, high EI values that certain arms were visited repeatedly at the expense of other arms not being explored. Boxplots represent median and interquartile range. Values for final R1 session (black background columns), EdRR 1, and EdRR 2. Certain outliers notwithstanding, the R1 population, who explored less than controls in the final R1 session, nevertheless explored the novel environment during EdRR 1 at almost optimal levels, with a lower median population EI even than controls. During EdRR 2, in the same and therefore, logically, now less novel environment, the R1 population, but not controls, had a significantly higher median EI score, which we will see corresponded with higher initial probability of choosing S1 (‘P(S1)’). **(B)** Linear regression analysis revealed no correlation between individual EI values and strength of within-error R1 bias, indicating independence of the cognitive mechanisms responsible for these behaviors. With respect to this, note in (A) that certain optimal explorers in EdRR 1 failed to fully explore all arms in EdRR 2. **(C)** Analyzing in more detail, we represented as histograms the trials in which animals made their first S0 choice (1 bin = 1 trial). This revealed that R1 trained animals were already choosing earlier than could be predicted from R1 performance levels in the final R1 session, but this leftward skewing increased significantly in EdRR 1 with 57.5% or R1 trained animals choosing S0 on the first trial. This proportion decreased in EdRR 2, though still skewed towards the beginning of the session. Controls first chose S0 according to a population probability closer matching random distribution (see main text for details). **(D)** Average population level P(S1) per trial for final R1, EdRR 1, and EdRR 2. This allows visualization of pronounced initial exploratory behavior in trials 1 + 2 of EdRR 1, followed by persistently high P(S1 values in remaining EdRR 1 trials and in EdRR 2.

#### 2.2 Decreasing novelty gives way to interference

To more rigorously evaluate the specific role of novelty in this exploratory behavior, we posited that its impact should be maximal upon first exposure to the novel environment (i.e. EdRR 1) then lessened in subsequent sessions. We therefore calculated individual EI values for both the previous (final R1) and subsequent (EdRR 2) sessions and compared these to the EdRR 1 values. Due to the intentionally accentuated qualitative environmental differences between the EdRR and prior R1 environments, direct within-individual comparison between final R1 and EdRR 1 are at best indicative (e.g. since R1 had no spatial landmarks, animals were less able to discern whether they had visited all S0 arms or simply some S0 arms many times; R1 uses all 8 arms of the maze, EdRR only 6, etc.). However, comparing final R1 session behavior (black columns in figure 3A) between R1 trained and control animals, we observed that 30/40 (75%) R1 trained animals did *not* fully explore the R1 environment, significantly more than the 6/19 (32%) in control animals (one-way ANOVA with pairwise Tukey HSD post-hoc between groups, with unbiased Cohen effect size: *F*(1, 57) = 6.12, *p* = 0.016, *d* = 0.69). Both groups subsequently displayed an increase in exploratory behavior when introduced to the novel EdRR environment. However, upon second exposure (EdRR 2) the median EI value significantly *increased* in the R1 but not the control population, reflecting an increase in the R1 population’s preference for choosing S1 (figure 3A; R1 population, 0.0625 in EdRR 1 vs 0.3125 in EdRR 2, paired t-test, *t*(39) = 2.6, *p* = 0.013; controls, 0.1875 in EdRR 1 vs 0.25 in EdRR 2, paired t-test, *t*(18) = 0.3, *p* = 0.76). The number of individual R1 trained animals who failed to explore all available arms also *increased* from 2/40 (5%) in EdRR 1 to 5/40 (12.5%) in EdRR 2, whereas in the control group no animals failed to explore all available arms in either EdRR 1 or 2. Yet strikingly, all R1 trained animals who *failed* to fully explore during EdRR 2 had been optimal or quasi-optimal explorers (i.e. EI < 0.1) in EdRR 1. Moreover, there was no correlation between R1 population EI values and level of within-error bias, either in EdRR 1 (figure 3B; *R*² = 0.063) or EdRR 2 (supplementary figure S.1D; *R*² = 0.007). These observations again suggest that distinct, albeit interacting, cognitive processes are responsible for each of these behaviors.

#### 2.3 Exploring space and exploring rules

We next identified the trial on which each animal made its first S0 choice, comparing again between the final R1 session, EdRR 1, and EdRR 2 (figure 3C). In the final R1 session, we can see that even here, despite having learnt that S0 is associated with reward *absence*, the majority of R1 trained animals made their first S0 (i.e. *exploratory*) choice towards the beginning of the session (figure 3C, top, red bars; 24 animals out of 40, i.e. 60% in the first, second, or third trials). In the case of control animals, we predicted that they would make their first S0 choice in the R1 environment according to a distribution commensurable with random choice behavior, and this is what we observed. All control animals chose S0 for the first time within the first five trials, following a distribution of, respectively, 10, 5, 1, 2, 1 mice per trial (figure 3C, top, blue bars). However, when initially introduced to the novel EdRR environment, the proportion of R1 trained animals choosing the S0 arm on the first trial jumped to 23/40 (57.5%) (figure 3C, middle, red bars). 15 of the remaining 17 R1 trained animals (15/40, 37.5%) chose S0 on the second trial, 1 on the third, and, finally, the last of the R1 trained animals on trial 8. Importantly, the timing of these initial EdRR environment S0 choices can be explained neither by strength of individual R1 performance in the final R1 session 24 hours earlier nor by a random choice behavior distribution. This indicates a specifically exploratory boost due primarily to environmental novelty *per se*. Recall that the very first presentation of each pair in each session of EdRR necessarily constitutes a *confirmation* of R1. In these three trials, S1 arm choices are rewarded, S0 arm choices unrewarded. Indeed, one reason we included this session-leading ‘anchor’ of R1 confirmations in our EdRR experimental design was to enable a certain level of separation between behavioral response to *environmental* novelty and behavioral response to *response-outcome* novelty. In EdRR 2, animals still tended to explore S0 surfaces early, but markedly *less* so than in EdRR 1 (figure 3C, bottom). The initial novelty of the EdRR environment thus appeared to increase exploratory behavior in two senses. First, ‘topologically’, with respect to physical exploration of all accessible regions of the environment. Second, ‘nomologically’ (from Greek *nomos*, meaning law, rule, custom), with respect to pointedly exploring beyond the bounds dictated by the R1 rule, an exploration which translates as an initial active preference for choosing S0 arms.

#### 2.4 Explore first, whatever the expense

Finally, we looked at the population level conditional probabilities of choosing the S1 arm (‘P(S1)’) in every trial of the first two EdRR sessions (with specific focus on the initial three trials; figure 3D, middle & bottom), comparing them to the P(S1) values from their final R1 training session (figure 3D, top). Here we saw that P(S1) in the R1 population on the first EdRR 1 trial was 0.425, in controls 0.368 (figure 3D, middle). We tested the statistical significance of these results against both the 0.5 chance value expected if a population were choosing randomly and also against both populations’ P(S1) values for the first trial of the final session of R1 training (Final R1, P(S1|1^st^ trial): R1 population = 0.825; controls = 0.474). These analyses revealed no significant difference compared with chance level in either population (t-tests with Welch correction against 0.5: R1 trained, *t*(39) = 0.95, *p* = 0.349; controls, *t*(18) = 1.16, *p* = 0.262) but a significant difference in the R1 population when compared with their first trial P(S1) in the final R1 session (t-tests with Welch correction: R1 trained, *t*(39) = 5.05, *p* < 0.0001; controls, *t*(18) = 0.93, *p* = 0.365). On the second trial, however, P(S1) decreased further to 0.175 for the R1 population, and to 0.263 for the control group. Importantly, this means that out of those R1 trained animals who chose to explore the S0 arm on the first trial (i.e. 23/40, who were therefore *not* rewarded on this trial and thereby experienced a confirmation of R1 ‘in the negative’, i.e. S0 + no-reward), a significant majority (18/23, 78.3%) nevertheless did not consequently choose the S1 arm on the second trial, but instead again chose to explore an S0 arm, foregoing the possibility of reward under the modalities of the R1 rule they had nevertheless internalized. In control animals also, 10/12 (83%) who chose S0 on the first EdRR 1 trial (and were thereby unrewarded) chose S0 again on the second trial. Out of those R1 trained animals who chose the S1 arm on the first trial, thereby receiving a reward and a ‘positive’ confirmation of R1 (i.e. S1 + reward), again the most significant proportion (15/17, 88.2%) nevertheless then chose to explore the S0 arm on the second trial. In control animals, 4/7 (57%) who had chosen S1 on the first trial chose S0 on the second trial, commensurate with both random choice behavior and their choice behavior in the final R1 session. This low second trial P(S1) value was significantly different from chance level in both the R1 population (*t*(39) = 5.34, *p* < 0.0001) and in controls (*t*(18) = 2.28, *p* = 0.035), but significantly different from P(S1) on the same trial in the final R1 session only in the R1 population (*t*(39) = 8.6, p < 0.0001; controls, *t*(18) = 2.03, *p* = 0.057). It was only on EdRR 1 trial 3 that population level decision behavior shifted in the S1 direction, in both the R1 population and controls. Even at the extremes of individual behavior, only 1 R1 trained animal chose to explore three consecutive (rewarded) S1 arms in EdRR 1, whereas 3 chose three consecutive (unrewarded) S0 arms. This serves to underline the cognitive implications of all the above results: under the initial effect of high novelty, the drive to explore can dominate even at the cost of repeated foregone reward.

During the rest of EdRR 1, P(S1) in the R1 population stayed above 0.5 on every trial except the final one (a trial of difficulty level 0), whereas in the control group P(S1) oscillated more or less evenly above and below 0.5 (see below). Compatible with our hypothesis of decreasing impact of novelty with repeated exposure to the novel environment, in EdRR 2 we did not observe the same low P(S1) values in the initial trials (figure 3D, bottom). Similar to EdRR 1, however, R1 population P(S1) values subsequently stayed above 0.5 throughout the remaining trials of EdRR 2. Indeed, in both EdRR 1 and EdRR 2, overall P(S1) values were significantly different to chance level in the R1 population (T-test with Welch correction and unbiased Cohen effect sizes: EdRR 1, *t*(22) = 5.05, *p* < 0.0001, *d* = 1.05; EdRR 2, *t*(22) = 7.42, *p* < 0.0001, *d* = 1.54; note that the effect size of this difference is ∼1.5 times greater in EdRR 2 compared to EdRR 1) but not in controls (EdRR 1, *t*(22) = 0.68, *p* = 0.5; EdRR 2, *t*(22) = 1.96, *p* = 0.06). It is also clear from figure 3D (bottom) that troughs in the R1 population P(S1) values preferentially occurred on trials of difficulty level 0, leading us onto our next focus of analysis.

### 3. R1 bias increases as a function of increasing EdM cognitive uncertainty

Based on recent analysis from our lab, we expected errors in the EdRR task to increase as a function of EdM trial difficulty, as a result of concomitant increase of retrieval-induced forgetting (RIF) of trial-relevant mnemonic episodes (Stevens et al., 2023). Furthermore, we predicted that the more EdM episodes were representationally weakened via RIF, the more interference from R1 would bias decision-making. To recall (see Materials & Methods), level 0 trials are most closely equivalent to basic spontaneous spatial alternation tasks, since the pair presented during such trials *n* is the same pair that was presented in the directly previous trial *n-1*, i.e. without any interposed presentations of other pairs which would provoke active forgetting of the *n-1* episode and concomitant RIF. In short, this interpretation advances that performance is highest on level 0 trials, not simply because they are more ‘recent’ in time, but because the mnemonic representation of *n-1* necessary for accurate response on *n* is significantly less likely to have been weakened via RIF (Stevens et al., 2023).

#### 3.1 EdM errors increase as a function of trial difficulty

In order to achieve stronger statistical power in our trial difficulty level analyses, EdM session performances were aggregated into blocks of three, according to an experimental design whereby each block is conceived to comprise exactly 12 trials at each level of difficulty (5 levels of trial difficulty = 60 trials in total per block, excluding initial presentation of each pair in each session). At this level of analysis, we observed that, as with naïve mice from previous studies (Stevens et al., 2023), the percentage of EdM choice errors in both the R1 and control populations increased as a function of trial difficulty (figure 4A; top, three-way ANOVA, major effect of ‘Difficulty’: *F*(4, 3500) = 130, *p* < 0.0001). Importantly, it is also clear from this graphical representation that only in block 1 (figure 4A, top left) and only on the most difficult trials, did the R1 population make significantly more errors than controls (pairwise t-tests with Bonferroni correction, between ‘Group’ difference: difficulty level 3, *t*(103) = 3.01, *p* = 0.016; difficulty level 4, *t*(105) = 3.02, *p* = 0.016). Errors made by the R1 population on difficulty level 3 trials in block 1 went significantly beyond chance level, i.e. the percentage of errors that could be explained by random choice behavior (i.e. ∼50%), and a similar but not significant trend was seen with errors made on level 4 trials (independent t-tests: difficulty level 3, *t*(119) = 4.29, *p* < 0.0001; difficulty level 4, *t*(119) = 1.61, *p* = 0.1).

**Figure 4.**
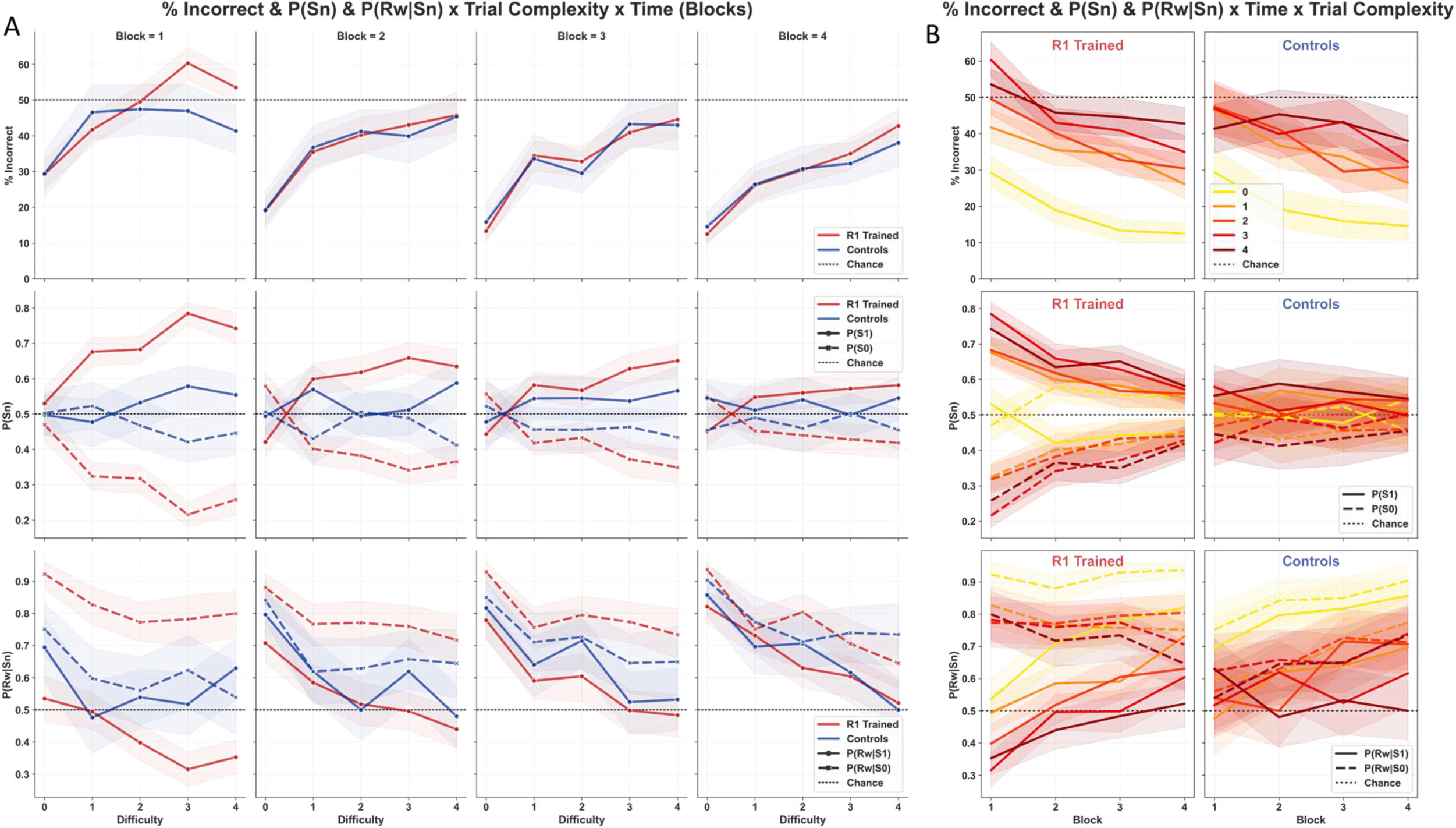
R1 interference in EdRR very significantly impacted by trial difficulty. By aggregating sessions into blocks of 3, each block contains exactly 12 trials of each trial difficulty level, allowing for more robust statistical analysis while still maintaining a sufficient dimension of evolution over time. Elsewhere, we analyze all trials, averaged across difficulty levels and including initial trials (see figure 5). Here we analyzed errors, probability of choosing S1 or S0 (‘P(Sn)’), and probability of being rewarded (outcome) given that S1 or S0 had been chosen (‘P(Rw|Sn)’), all as a function of trial difficulty. We provide two visualizations of the same findings. In (A), trial difficulty from 0 to 4 is represented on the x-axis and each column represents a block of EdRR, from 1 to 4. In (B), EdRR blocks are represented on the x-axis, values according to trial difficulty are represented as color-coded curves, from yellow (level 0) to burgundy (level 4), and each column represents one of the populations: R1 trained, left; controls, right. Error bands in all cases represent 95% confidence intervals. **(A)** + **(B)**, top row: % of EdM errors was significantly impacted by trial difficulty, matching what we expected based on the existing classical EdM task literature. Level 0 performance in both groups was particularly strong, even in block 1. By contrast, R1 trained animals performed significantly worse than controls only on level 3 + 4 trials and only in block 1. Their performance in these trials was below what could be explained by chance. In subsequent blocks, EdM performance in both groups improved as a function of time, nevertheless always as a function of trial difficulty. **(A)** + **(B)**, middle row: P(Sn) expressed as a probability between 0 and 1. P(S1) increased highly significantly as a function of trial difficulty in R1 trained animals. This dependence of P(Sn) on trial difficulty decreased gradually but was still visible and significant in all blocks. P(Sn) on level 0 trials thus demonstrated the least impact of R1 training. Indeed, a reflection of high EdM performance on level 0 trials is precisely that R1 trained animals were capable of actually choosing S0 more often than S1, but on these easiest trials only (more details in text). In controls, by contrast, P(Sn) fluctuated on all trial difficulty levels except level 4 where a slightly but consistently higher P(S1) value was observed, potentially indicating a consistent response strategy involving surface on these trials. **(A)** + **(B)**, bottom row: P(Rw|Sn) expressed as a probability between 0 and 1. The probability of R1 trained animals being rewarded was significantly higher on trials of all difficulty levels when they chose S0 as opposed to S1. This is a reflection of the fact that the majority of their EdM errors were committed on S1 arms, implying that reward was most often ‘waiting’ for them on the S0 arm of a pair. P(Rw|S0) did not significantly evolve over time, except on level 4 trials, where it decreased slightly. Rather, with repeated EdRR training, P(Rw|S1) gradually rose up to almost comparable levels as P(Rw|S0) on all difficulty levels, though even this improvement with time had a trial difficulty dependence most easily visualized in block 3 where P(Rw|S1) decreases step like from 0.8 at level 0, to 0.6 on levels 1-2, to 0.5 on levels 3-4. Whereas, in contrast, P(Rw|S0) is higher on level 0, as expected, but almost equal at ∼0.75 on all trial difficulty levels.

We also observed that the number of errors decreased significantly and consistently across all difficulty levels as a function of time/repeated training (three-way ANOVA, major effect of ‘Block’: *F*(3, 3500) = 72, *p* < 0.0001). In figure 4B (top) we have provided an alternative graphical representation of these same evolutions to facilitate understanding of the evolution in time of performance at each difficulty level. Importantly, this representation also renders even clearer just how *categorically* fewer errors occurred on level 0 trials (yellow lines), across all training blocks and in both the R1 and control populations (pairwise t-test between % errors performed at trial difficulty level 0 and level 1, averaged across groups and blocks: *t*(58) = 5.47, *p* < 0.0001).

#### 3.2 S1 choice probability increases as a function of trial difficulty

In figure 2C above, we measured R1 interference in EdM performance as a within-error R1 bias index. This demonstrated that although R1 animals and control animals made comparable overall numbers of errors, the nature or quality of these errors was significantly and persistently biased towards S1 in the R1 population. However, such an analysis presented alone may transmit the misleading idea that bias exists only when mistakes are made. In fact, bias induced by previous learning cannot and should not be reduced to something that manifests only when errors arise, precisely because error *outcomes* are as much a function of the external environment as they are of an organism’s internal neurocognitive processing. Where there is bias in cognition, we should expect it to manifest independently of whether a given subject-environment interaction gives rise to a correct or to an incorrect outcome (see Discussion below). For this reason, we also measured the overall trial-by-trial probability of choosing S1 or S0, independently of whether or not that choice led to an EdM error (figure 4A & figure 4B, middle rows).

In a behavioral paradigm based on spatial alternation in which the space in question is covered half and half with two different surfaces, all other things being equal besides, we should expect the probability of exploring each surface type (P(Sn)) to converge towards 0.5 (with a certain calculable margin of variance), whether this be in animals performing the EdM task perfectly or purely randomly. Comparing this to our results from each block (figure 4A-B, middle row), what we observed in control animals is that the mean choice probability of exploring each surface (P(S1) and P(S0)) lay between maximal extremes of 0.6 and 0.4, respectively, across all blocks and across all levels of trial difficulty. As a result, the mean probability of choosing one surface over another achieved statistical significance at only three discrete points: level 3 trials in block 1 (pairwise t-test with Bonferroni correction: *t*(112) = 3.42, *p* = 0.004), and levels 1 and 4 trials in block 2 (pairwise t-tests with Bonferroni correction: *t*(112) = 2.91, *p* = 0.02; *t*(112) = 3.61, *p* = 0.002, respectively). This result indicated a dependent interaction between trial difficulty and surface choice in the control group which emerged as statistically significant on difficulty levels 3 and 4 only when averaged across all blocks (pairwise t-tests with Bonferroni correction, interaction ‘Difficulty*P(Sn)’: level 3, *t*(454) = 2.74, *p* = 0.031; level 4, *t*(454) = 5.33, *p* < 0.0001, respectively). In sum, a significant global effect of surface on choice was observed in the control group with a significant interaction between difficulty and surface (two-way ANOVA averaged across all blocks: *F*(1, 2270) = 34, *p* = 0.001, with Tukey HSD post-hoc; *F*(4, 2270) = 3.42, *p* = 0.009, respectively).

Despite this slight but significant surface effect in controls which could not be explained by prior learning (analyzed further below), P(Sn) was nevertheless significantly more impacted in the R1 population (two-way ANOVA, interaction ‘Group*P(Sn)’: *F*(1, 7076) = 104, *p* < 0.0001). Furthermore, across all blocks, P(S1) in the R1 population increased very significantly (mirrored by a proportional *decrease* in P(S0)) as a function of trial difficulty (pairwise t-tests with Bonferroni correction: *t*(958) > 4, *p* < 0.0001 for all levels; unbiased Cohen effect sizes, level 0, *d* = 0.3; level 1, *d* = 0.85; level 2, *d* = 0.86; level 3, *d* = 1.31; level 4, *d* = 1.15), though the magnitude of the difference did decrease gradually as a function of time/repeated training (two-way ANOVA, interaction ‘P(Sn)*Block’: *F*(3, 4792) = 69, *p* < 0.0001; ANOVA with Tukey post-hoc on P(Sn) in block 4: *F*(1, 1198) = 35.2, *p* = 0.001; unbiased Cohen effect size, *d* = 0.34). In fact, in the visual comparison seen in supplementary figure S.2E, we can see that, during block 1 of EdRR, P(S1) on trials of difficulty level 3 and 4 (designated ‘hard’ in the graph) was almost as high as it had been in the final R1 session, despite the fact, just seen, that a clear majority of P(S1) choices in EdRR block 1 were giving rise to unrewarded errors, i.e. R1 *dis*confirmations.

P(S1) and P(S0) values were most closely matched in the R1 population on difficulty level 0 trials in block 1 (mean values P(S1) = 0.53, P(S0) = 0.47). In subsequent blocks, P(S0) increased to be significantly higher than P(S1) on level 0 trials only (this finding is highlighted by the graphical ‘mirror’ representation in the middle rows of figure 4A-B, intended to make clear where P(S0) becomes greater than P(S1)). Although this S0 preference at level 0 may seem counter-intuitive, it is easily explained by the fact that the same animals were significantly more likely to choose S1 arms at all other trial difficulty levels, meaning that whenever a level 0 trial *n* occurred there was a statistically higher probability that the animal had chosen the S1 arm on the immediately previous *n-1* presentation of the same arm pair. Thus, if the animal spatially alternated on a level 0 trial (which, referring to figure 4A, top, we can see that the R1 population was doing on average 80% of the time in blocks 2 to 4), this would necessarily imply choosing the S0 arm significantly more often than the S1 arm on level 0 trials. Still, that R1 trained animals retained sufficient executive control to indeed choose S0 more often than S1 on level 0 trials is note-worthy in itself and corroborates our previous active forgetting interpretation of EdM performance (Stevens et al., 2023). Notably, this interpretation entails that RIF representationally weakens actively forgotten episodes. But since, on level 0 trials, the *n-1* memory episode required for accurate response on trial *n* is unlikely to have undergone RIF, it is therefore representationally stronger and putatively more resistant to intrusive interference, e.g. from prior R1 learning.

### 4. R1 bias impedes updating of S1 choice behavior

Next, we analyzed the resultant conditional probabilities for animals from both groups to obtain a food reward (correct EdM response) given the surface chosen (P(Rw|Sn)), again also as a function of trial difficulty (figure 4A-B, bottom rows). Note that, in contrast to P(Sn), where both perfect and purely random EdM performances would result in P(S1) and P(S0) values converging towards 0.5, a hypothetical perfect EdM performance in the EdRR environment would result in P(Rw|S1) = 1 and P(Rw|S0) = 1, while a purely random performance would result in P(Rw|S1) and P(Rw|S0) values converging towards 0.5. Tracking P(Rw|Sn) values was an essential first step towards understanding how P(Sn) values were updated in each group over the course of repeated EdRR sessions.

#### 4.1 Significant impact of R1 bias on reward probability given surface choice

A general trend for the probability of reward on both surfaces to increase across time/repeated training was observed, with no significant difference between the two groups in overall P(Rw) (one-way ANOVA: *F*(1, 6473) = 0.38, *p* = 0.54; this is a logical implication of overall EdM performances being similar between the two groups, as seen in figure 3A above). There was, however, a highly significant interaction between ‘Group’ and ‘P(Rw|Sn)’ (two-way ANOVA with Tukey HSD post-hoc: ‘Group*P(Rw|Sn)’, *F*(1, 6471) = 82, *p* = 0.001). In the R1 population, P(Rw|S1) was very significantly lower than P(Rw|S0) (figure 4A-B, bottom; one-way ANOVA with pairwise Tukey HSD post-hoc: *F*(1, 4340) = 529, *p* = 0.001, Cohen effect size, *d* = 0.699), an expected result given that their P(S1) was significantly higher than their P(S0). This significant difference between P(Rw|S0) and P(Rw|S1) values in the R1 population was observed at all trial difficulty levels across blocks 1 to 3 and persisted, albeit less significantly, in block 4 also (pairwise t-tests with Bonferroni correction averaged across all blocks: ‘P(Rw|Sn)*Difficulty’, *p* < 0.0001 at all difficulty levels). In controls too, a trend for P(Rw|S0) to be higher than P(Rw|S1) emerged starting in block 2, but reached statistical significance only in block 4 on difficulty level 4 trials (pairwise t-tests with Bonferroni correction averaged across all blocks, interaction ‘P(Rw|Sn)*Difficulty’: levels 0-2, *p* > 0.9; level 3, *p* = 0.34; level 4, t(106) = 3.3, *p* = 0.007). Importantly, however, we will now see that, in controls but not in the R1 population, global surface-based differences in P(Rw|Sn) and P(Sn) were commensurate with each other (figure 5A-B).

**Figure 5.**
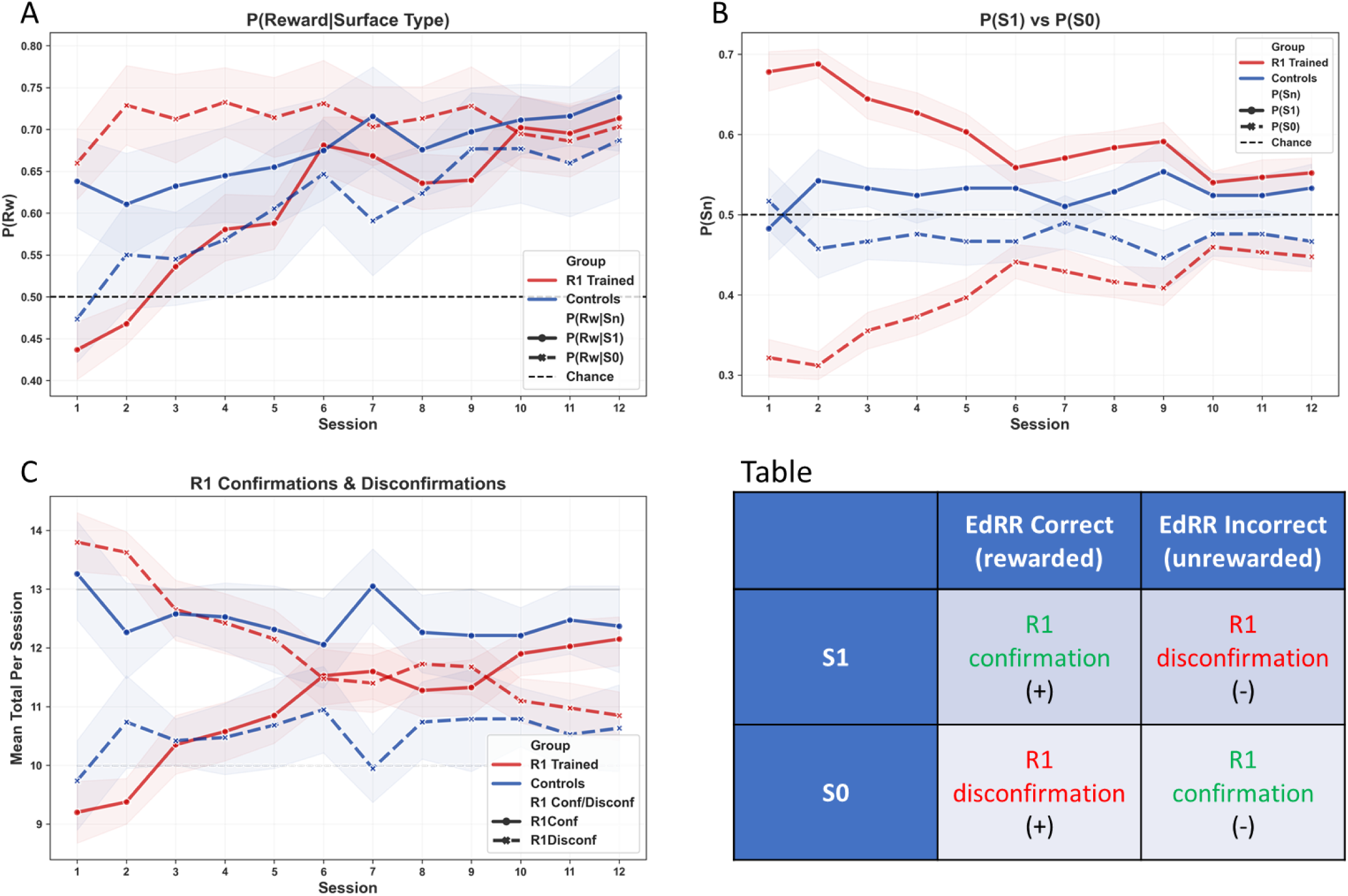
Controls but not R1 trained animals update P(Sn) in accordance with P(Reward|Surface Type). Overall per session P(Rw|Sn) values, averaged across all difficulty levels and encompassing initial per session trials on each pair, in which P(Rw|S1) = 1 and P(Rw|S0) = 0. P(Sn) per session is represented beside this, allowing easy visualization of the contrast between the R1 trained and control groups with respect to the relationship between, specifically, P(Rw|S1) and P(S1). All error bands represent 95% confidence intervals (for detailed statistical analyses, see main text). **(A)** P(Rw|Sn). The R1 trained and control populations experienced opposing outcome values in this regard. In R1 trained animals, P(Rw|S0) was higher than P(Rw|S1) throughout the first 3 blocks. Their P(Rw|S0) did not significantly evolve over repeated training, rather P(Rw|S1) gradually increased over time until the two values came level in block 4 only. By contrast, in controls, P(Rw|S1) was slightly but robustly higher than P(Rw|S0) across all 12 sessions of EdRR, a contribution to which comes from the initial per session trials in which P(Rw|S1) = 1. **(B)** P(Sn). The fact that the control group displayed a robust trend, beginning only from EdRR 2, to slightly favor S1 thus seems to find at least partial explanation in this group’s robustly higher P(Rw|S1) values. It is also clear to what extent P(S1) values in the R1 population remained persistently resistant to the fact that they were being rewarded less often when they chose S1, and precisely more often when they chose S0. **(C)** + **(Table)** Mean total R1 confirmations and disconfirmations per session per group. Our EdRR protocol was designed specifically in such a way that trial-by-trial outcomes could be qualified as either confirmations or disconfirmations of the previously learned R1 rule. In the Table, we recap briefly which EdRR outcomes constitute an R1 confirmation and which a disconfirmation. Recalling that in EdRR the outcome of the initial per session trial on each pair is always a R1 confirmation, this implies that both hypothetical perfect and perfectly random EdRR performances will generate a total number of R1 confirmation outcome trials converging towards 13 (out of 23) and of R1 disconfirmation outcome trials converging towards 10. The horizontal unbroken and dashed light grey lines at 13 and 10, respectively, are visual anchors to help represent this. Thus, it is visually clear that total numbers of confirmations and disconfirmations in control animals did tend to converge towards these modelled values, whereas in the R1 population initial values were inverted in this respect; many more R1 disconfirmations being the result of many incorrect (in terms of EdM) S1 choices. Only during block 4 did this population begin to converge towards modelled values. Referring back to (B), the visible lack of decrease in P(S1) between EdRR 1 and EdRR 2 in the R1 population is all the more striking in light of the number of R1 disconfirmations we see they had experienced during EdRR 1.

It was essential to also analyze *global* P(Sn) and P(Rw|Sn), notably including the initial three trials from each session which are *not* included in analyses by EdM trial difficulty (since, as sampling trials, they have no difficulty level) and wherein, necessarily, P(Rw|S1) = 1 and P(Rw|S0) = 0. The response-outcomes of these initial trials would therefore exert an important influence on each animal’s *cumulative* per session experiences of P(Rw|Sn). What we observed here is that when the initial trial P(Rw|Sn) values were included, P(Rw|S1) and P(Rw|S0) were drawn closer together in the R1 population than they had been when analyzed only according to EdM trial difficulty, coming into almost perfect alignment by the final three sessions/block 4 (figure 5A). In controls, interestingly, inclusion of the initial trials revealed that their P(Rw|S1) values were globally higher than P(Rw|S0) across all EdRR sessions (figure 5A), a fact which was hidden when these values were analyzed according to EdM trial difficulty only. Hence, looking at figure 5A and 5B as we have presented them here side-by-side, we are invited to entertain the possibility that the trend observed in controls for P(S1) values to rise slightly higher than P(S0) values over time may be at least in part explained as a cognitive behavioral adjustment (e.g. some kind of putatively Bayesian updating) to this population’s consistently higher P(Rw|S1) values. Lending weight to this hypothesis of an ongoing P(Rw|Sn) updating in controls is the observation that P(Rw|S1) was already higher than P(Rw|S0) in EdRR 1, whereas P(S1) rose above P(S0) only starting from EdRR 2 (figure 5A-B, see Discussion).

In supplementary figure S.2A, for completeness, we have also isolated and represented the session-by-session P(S1) vs P(S0) values for these initial trials only, where we can again see the dramatic change in P(Sn) between the high novelty EdRR 1 and subsequent lower novelty EdRR sessions in R1 trained animals.

#### 4.2 Lag in S1 update despite R1 disconfirmations outweighing R1 confirmations

In EdRR task conditions, R1 *confirmations* can take two forms: rewarded EdM outcome on an S1 arm, or unrewarded outcome on an S0 arm. Likewise, R1 *disconfirmations* can also take two forms: rewarded EdM outcome on an S0 arm, or unrewarded outcome on an S1 arm (see figure 5, table). Recall also that on initial per session presentation of each pair, arm choices can be neither correct nor incorrect in terms of the classical EdM task (based on making a choice on trial *n* according to the choice made on the same pair on trial *n-1*, thus excluding the initial *n* since it has no corresponding *n-1*). In EdRR, upon initial per session presentation of each pair, only S1 arms were rewarded, meaning that regardless of whether S1 or S0 was chosen, response on these initial three trials *always* gave rise to an R1 confirmation (see Materials & Methods).

In figure 5, we present the all-trial inclusive, session-by-session, trial difficulty independent representations of P(Rw|Sn) and P(Sn) side-by-side in order to make the relations between them clearer. Hence, in figure 5A-B, in the R1 population but not in controls, we can observe a lag in rule-revision manifest as a delay in P(Sn) being updated as a function of P(Rw|Sn). We have similarly included a representation of the average session-by-session number of R1 confirmations and disconfirmations experienced by the two populations during EdRR (figure 5C). During the first five EdRR sessions, we can thereby see that the R1 and control populations had diametrically opposed experiences in terms of R1 confirmations and disconfirmations. The R1 population experienced significantly more disconfirmations than confirmations, while the controls experienced the opposite.

In order to better frame these results, two additional elements were included in figure 5C. Since the initial per session trials on each pair necessarily constitute R1 confirmations, this entails that, during EdRR, both a perfect EdM performance and a purely random EdM performance will give rise to a per session number of R1 confirmations converging towards 13 (solid light grey horizontal line at y = 13) while R1 disconfirmations will correspondingly converge towards 10 (dashed light grey horizontal line at y = 10). Visually, these elements allow us to see that, in controls, R1 confirmation and disconfirmation averages remained close to these modelled values throughout EdRR, whereas in the R1 population, in the early sessions, the average total R1 confirmation and disconfirmation values were inverted with respect to the same modelled values (mean values EdRR 1, R1 confirmations = 9.2; R1 disconfirmations = 13.8). Across the 12 EdRR sessions, observed values came close to modelled values in the R1 population only during block 4, but even then still not as close as in controls. Thus, while both the R1 and control populations’ R1 confirmation and disconfirmation values were significantly different to the modelled values, this difference was significantly greater in the R1 population (t-tests with unbiased Cohen effect sizes: R1 trained, *t*(479) = 24.6, *p* < 0.0001, *d* = 1.12; controls, *t*(227) = 5.2, *p* < 0.0001, *d* = 0.34; one-way ANOVA with Tukey HSD post-hoc between ‘Group’: *F*(1, 706) = 112.5, *p* = 0.001).

Finally, it is of particular interest in terms of rule revision to note that there was no decrease in P(S1) between sessions 1 and 2 in R1 trained animals (figure 5B; pairwise t-test: *t*(39) = 0.56, *p* = 0.5). This relative *absence* of revision of P(S1) observable in EdRR 2 can here be further considered in light of the average 13.8 R1 disconfirmations experienced during EdRR 1, accounting for 60% of all trials.

### 5. R1 bias persists longest in deliberative behaviors

#### 5.1 Population differences in decision latency fade during EdRR

In the latter half of the R1 training phase, prior to introduction into the EdRR environment, the R1 tactile-discrimination population displayed significantly higher overall decision latency values compared to controls (recall: controls rewarded regardless of surface chosen), and also significantly higher within-group decision latency values on rewarded S1 compared to non-rewarded S0 arms (data not shown). We interpreted these observations as a behavioral marker of the cognitive effort required to inhibit the ‘innate’ exploratory drive in order to maintain a tactile ‘win-stay’ strategy (Stevens et al., forthcoming). Upon subsequent introduction to the EdRR environment, we observed an initial putatively novelty-induced increase in overall decision latency in both the R1 trained and control populations when compared to their final R1 session values (figure 6A; R1 trained, ∼2900 ms up from ∼2500 ms; controls, ∼2600 ms up from ∼2000 ms). However, decision latency remained significantly higher in the R1 population only during EdRR 1, after which there was no longer any significant difference in overall decision latency between the two populations (pairwise t-tests with Bonferroni correction and unbiased Cohen effect sizes: EdRR 1, *t*(115) = 3.38, *p* = 0.01, *d* = 0.54; EdRR 2, *t*(92) = 2, *p* = 0.64). Furthermore, that decision latencies in R1 animals decreased steadily over repeated EdRR training, quickly dropping below mean final R1 phase decision latencies, also indicates that the spatial alternation ‘win-shift’ behavior elicited by the EdM protocol comes more spontaneously to mice than the tactile ‘win-stay’ S1 exploitation behavior required for initial R1 training. However, averaged across the three EdRR sessions of block 1, the R1 population did display higher decision latencies when their decision-making led them to explore S0 arms, which is to say, specifically during the process of deciding to ‘transgress’ the previously acquired R1 rule (figure 6A; difference approached statistical significance, pairwise t-tests with Bonferroni correction and unbiased Cohen effect sizes: *t*(238) = 2.5, *p* = 0.055, *d* = 0.3). These results can be interpreted as reflecting initial increase of cognitive conflict entailed by having to inhibit R1 in order to explore the novel EdRR environment. Importantly, however, this difference faded rapidly as a function of habituation to the spontaneous behavior-based EdM protocol.

**Figure 6.**
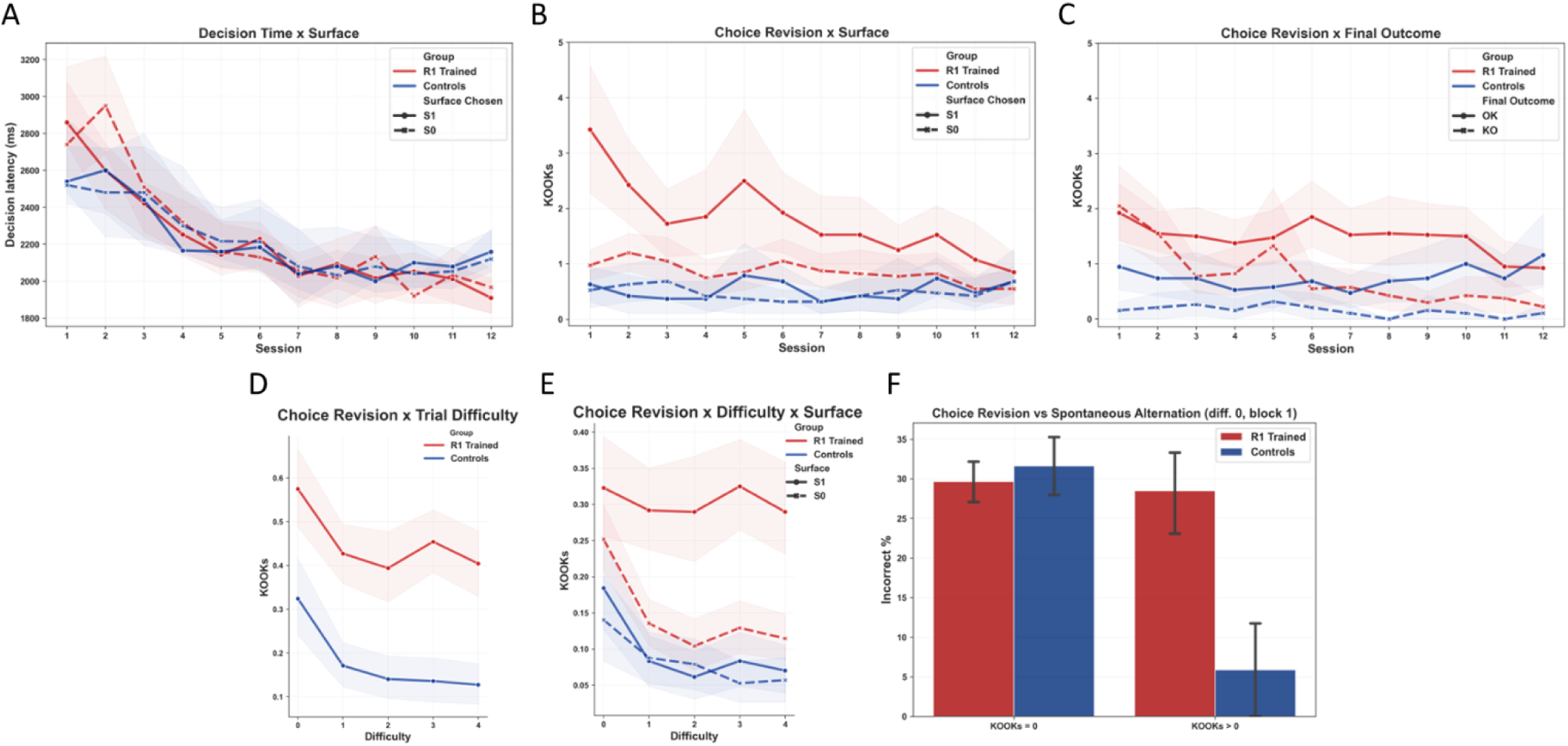
Deliberative choice revision behavior highly biased towards priorly acquired rule of action. To allow deeper investigation of pre-definitive choice deliberative processes, we developed analyses for measuring decision latency and choice revision behaviors. These behaviors proved to be highly revelatory with respect to R1 interference during EdRR and offered powerful insights into the cognitive and putative neurobiological mechanisms underpinning the cognition of rule revision. All error bands represent 95% confidence intervals (see main text for detailed statistical analyses). **(A)** Median decision latencies (in milliseconds) by surface and by session. Both populations displayed a significant time-dependent decrease in decision latency as a function of repeated EdRR training. In EdRR 1 + 2, R1 trained animals displayed a trend for significantly higher decision latency, particularly on S0 choice trials in EdRR 2, putatively an indication of cognitive effort to inhibit the R1 drive to choose S1. **(B)** Mean total choice revision by surface by session. R1 trained animals displayed around twice as much choice revision behavior as controls, with a significant tendency for the final decision to be an S1 choice. This was especially true in early EdRR sessions, decreasing gradually as a function of time, but with a persistent trend still even in EdRR 12. **(C)** Mean total choice revision by outcome by session. Controls were significantly more likely to engage in rectifying rather than error-inducing choice revision across all EdRR sessions. Choice revision in R1 trained animals in EdRR 1 + 2 was as likely to lead to error as to a correct response. Only from EdRR 6 onwards was choice revision in R1 trained animals significantly more likely to be rectifying. **(D)** Mean total choice revision by difficulty. Averaged across all blocks, R1 trained animals displayed significantly more choice revision than controls at all trial complexities. However, both groups engaged in significantly more choice revision behavior on level 0 trials. **(E)** Mean total choice revision by difficulty by surface. Dividing mean choice revision values according to surface further revealed that choice revision which terminated on S1 surfaces was equally high on all trial complexity levels, or rather was just as high on level 1-4 trials as it was on level 0 trials, in R1 trained animals but not controls. This indicates that striatal R1 cognitive content has a greater capacity to remain active and potentially intrusive at any trial complexity level, in contrast to cortico-hippocampal EdM content which putatively decreases in representational strength as an effect of active forgetting, thus leaving fewer EdM elements capable of provoking choice revision as trial difficulty level increases. **(F)** In level 0 trials, almost all choice revision in controls was rectifying whereas R1 trained animals were as likely to make an error with as without choice revision. This was driven by R1 bias (see main text).

#### 5.2 R1 bias manifest in increased choice revision deliberation

Next, we measured and analyzed physical choice revision behavior in both populations. Similar to classical “vicarious trial and error” (VTE) linked in the literature to representational accounts of predictive decision-making (Tolman, 1939; Redish, 2016), in our experimental design this behavior occurs when animals make an initial crossing of the threshold into one arm of a pair, only to revise their choice, return to the central platform, then choose again (for full details of differences between “choice revision” here and classical VTE, see Stevens et al., 2023). Strikingly, the R1 population engaged in significantly more overall choice revision than control animals (figures 6B-E, one-way ANOVA with Tukey HSD post-hoc: *F*(1, 1414) = 72.1, *p* = 0.001). We also observed that, as previously seen in classical EdM (Stevens et al., 2023), both populations performed significantly more choice revision on difficulty level 0 trials compared to all other difficulty levels (figure 6D; results averaged across all blocks; pairwise t-tests within ‘Difficulty’ between trial difficulty level 0 and each other level: *t*(58) > 3.9, *p*-values < 0.0002; among all other levels: *t*(58) < 1.4, *p*-values > 0.17). In (Stevens et al., 2023) we advanced that this trial difficulty-dependent *decrease* in deliberative behaviors is also a consequence of RIF-induced representational weakening, leaving the organism with less representational ‘stuff’ to deliberate with. In stark contrast, however, when we looked specifically at choice revision towards S1 arms we observed that, in contrast to classical EdM conditions, this was just as high on difficulty level 1 to 4 as on difficulty level 0 trials in the R1 population only (figure 6E). This indicates that R1 interference, putatively boosted by perception of R1 elements in the environment (i.e. haptic perception of surface types), does not so much override deliberation as re-fuel it, potentially by eliciting representational pattern completion (Horner & Burgess, 2014) which adds substance, albeit in an ultimately R1-biasing manner, back to the representational deliberative ‘stuff’ lost via RIF.

Corroborating this interpretation, choice revision decreased significantly over repeated EdRR training only in the R1 population, who correspondingly started from a significantly higher level than controls in EdRR 1 (figure 6B, repeated measures ANOVA within ‘Session’, R1 trained group: *F*(11, 429) = 9.2, corrected *p* < 0.0001; control group, *F*(11, 198) = 0.7, *p* = 0.74). In control animals, choice revision behavior was equally balanced between the S1 and S0 surfaces but did occur more often in a choice rectifying than in an error-inducing manner (figure 6B-C, one-way ANOVA with pairwise Tukey HSD post-hoc; within control group, between correct/incorrect outcome: *F*(1, 454) = 68.8, *p* = 0.001), again replicating previous analyses in naïve animals performing EdM (Stevens et al., 2023).

The R1 population, by contrast, revised their choice significantly more often towards S1 than towards S0 arms (one-way ANOVA with Tukey HSD post-hoc, within R1 trained group, between ‘Surface’: *F*(1, 958) = 56.1, *p* = 0.001). This effect was especially significant in EdRR 1 (figure 6E, pairwise t-tests with Bonferroni correction and unbiased Cohen effect sizes: EdRR 1, *t*(78) = 4.1, *p* = 0.001, *d* = 0.91; difference in EdRR 2, which appears significant on graph, was not significant following Bonferroni correction: *t*(78) = 2.7, *p* = 0.09). However, as seen in figure 6F, in EdRR 1 and 2, choice revision in the R1 population was equally likely to lead to an error as to a correction. Only from EdRR 6 onward was significantly more choice revision rectifying rather than error-inducing in the R1 population (pairwise t-tests with Bonferroni correction and unbiased Cohen effect sizes: EdRRs 1-5 + 11, *t*(78) < 2.5, *p* > 0.05; EdRRs 6-10 + 12, *t*(78) > 3, *p* < 0.05), but even during these sessions the overall significant trend for more choice revision to terminate with an S1 choice persisted (figure 6E, one-way ANOVA with Tukey HSD post-hoc on EdRRs 6 – 12: *F*(1, 558) = 20.1, *p* = 0.001). This trend did not switch at any point during the 12 sessions of EdRR.

Finally, since we had already observed that significantly more choice revision occurred on difficulty level 0 trials (supplementary figure 6D) and that EdM performance was significantly higher at difficulty level 0 trials (figure 4A, top), we hypothesized that there would be an observable interaction in the data between these two results. Hence, we isolated level 0 trial performances and separated them into two sets; those where no choice revision occurred (‘KOOKs = 0’) and those where at least one choice revision occurred (‘KOOKs > 0’) (figure 6F). During block 1, the R1 population engaged in choice revision behavior on 81 out of 480 (16.9%) level 0 trials, and controls on 21 out of 228 (9%). What we observed (figure 6F) is that the EdM performance of the two populations was similar on level 0 trials when *no* choice revision occurred, but that only controls committed significantly fewer EdM errors on those level 0 trials where they engaged in choice revision behavior, in contrast to the R1 population who were just as likely to make an error with or without choice revision (one-way ANOVA with Tukey HSD post-hoc, within each population between ‘KOOKs = 0’ and ‘KOOKs > 0’: R1 population, *F*(1, 173) = 0.05, *p* = 0.83; control population, *F*(1, 72) = 13.1, *p* = 0.001). To be precise, in the case of control animals, out of 21 trials on which choice revision occurred, only 1 of these gave rise to an error. With respect to surface, in the R1 population, 87% of the errors (102/117) made during block 1 on level 0 trials with no choice revision behavior occurred on S1, compared to 51% (35/68) in controls. On trials *with* choice revision, 100% of the errors (26/26) committed by the R1 population occurred on S1. As mentioned, in control animals only 1 error arose at level 0 *with* choice revision, precluding any meaningful analysis with respect to distribution of errors by surface. These population differences in choice revision behavior were most marked in block 1, as we had expected, but did also hold when analyzed across all training blocks averaged together, though with repeated EdRR training the R1 population also began to make less error-inducing choice revision (supplementary figure S.3A, one-way ANOVA with Tukey HSD post-hoc, within each population between ‘KOOKs = 0’ and ‘KOOKs > 0’: R1 population, *F*(1, 661) = 3.8, *p* = 0.051; control population, *F*(1, 282) = 14.4, *p* = 0.001). Averaged across all blocks, 236/316 (75%) of all level 0 errors *without* choice revision in the R1 population were S1 choices, while 38/43 (88%) of all level 0 errors *with* choice revision were. Taken together, these analyses confirmed what we had predicted: that increased cognitive deliberation was just as if not more likely to give rise to R1-biased choice behavior than choices made with less deliberation.

#### 5.3 Persistent R1 bias manifest as post-choice run time hesitation

Next, we looked at post-choice run times, the time taken to travel from the threshold of the definitively chosen arm (i.e. following any choice revision behavior which may have occurred) to the distal, reward distributor containing zone. The post-choice run time parameter arises from the physical design of the radial maze apparatus and is another central feature enabling analysis of deliberative processes as animals physically realize their choice, something that is not possible where response execution is quasi-instantaneous (e.g. lever press). Whereas in the latter half of R1 training, overall run time was higher in the R1 population (putatively due to control animals attaining higher *confidence* in being rewarded on every trial and therefore manifesting less post-choice hesitation), in the EdRR task we observed that overall run time, averaged across all sessions and surfaces, was actually slightly but significantly lower in the R1 population (figure 7A, one-way ANOVA with Tukey HSD post-hoc: *F*(1, 705) = 10.5, *p* = 0.001). Looking at the constitution of these overall values, we observed that, during the first five EdRR sessions, run times were significantly higher in R1 trained animals on S0 arms compared both to control animals (figure 7B, see 95% CI error bands; this difference translated into a statistically significant overall between-group difference in S0 run times; one-way ANOVA with Tukey HSD post-hoc: *F*(1, 705) = 7.6, *p* = 0.006) and to their own S1 arm run times (one-way ANOVA with Tukey HSD post-hoc: *F*(1, 956) = 77.6, *p* = 0.001: no significant difference between S1 and S0 run times in controls; *F*(1, 454) = 0.2, *p* = 0.67). In fact, this between-surface within-R1 population difference was significant even when we analyzed only blocks 3-4, i.e. even when there was no longer any significant session-by-session difference (one-way ANOVA with Tukey HSD post-hoc: *F*(1, 478) = 6.6, *p* = 0.001). Taking all this together, it seems that lower run times in the R1 population specifically on S1 arms drove this group’s overall, surface-independent lower run time values. Interestingly, when we looked at these group level differences in run times as a function of trial difficulty and repeated training (figure 7C), we observed that, in block 1 especially, run times were significantly greater in the R1 population on S0 arms at all trial difficulty levels, including level 0, on which, as we have seen, these animals were nevertheless performing relatively well (pairwise t-tests ‘Difficulty*Surface’ with Bonferroni correction and unbiased Cohen effect sizes, within block 1: level 0, *t*(174) = 5.3, *p* < 0.001, *d* = 0.72; level 1, *t*(155) = 3.4, *p* < 0.001, *d* = 0.49; level 2, *t*(117) = 5.5 *p* < 0.001, *d* = 0.81; level 3, *t*(81) = 3.1, *p* < 0.001, *d* = 0.55; level 4, *t*(82) = 4.2, *p* < 0.001, *d* = 0.76). We can also see from figure 6D just how reliable, across trial difficulty and repeated training, the trend for lower S1 run times was in the R1 trained group. For perspective, we can compare this surface choice breakdown of run times to an EdM outcome (correct or incorrect) breakdown (supplementary figure S.3B, bottom). Here, we can see that with repeated training, run times on incorrect response trials increase more steadily at all trial difficulty levels, but especially level 0, in the control group compared to the R1 trained group, i.e. towards a robust phenotype we also see in the classical EdM task for animals to have higher run times on incorrect response trials, putatively due to more pronounced lack of confidence in their reward-location prediction (Stevens et al., 2023). A suggestion of these run time results is that, in R1 trained animals, there is some kind of highly persistent, putatively partially affective (McDonald et al., 2004; McDonald & Hong, 2004) *under*-confidence in being rewarded on S0 and *over*-confidence in being rewarded on S1, even after extensive EdRR training throughout the majority of which P(Rw|S1) is in reality significantly lower than P(Rw|S0) (figure 5A).

**Figure 7.**
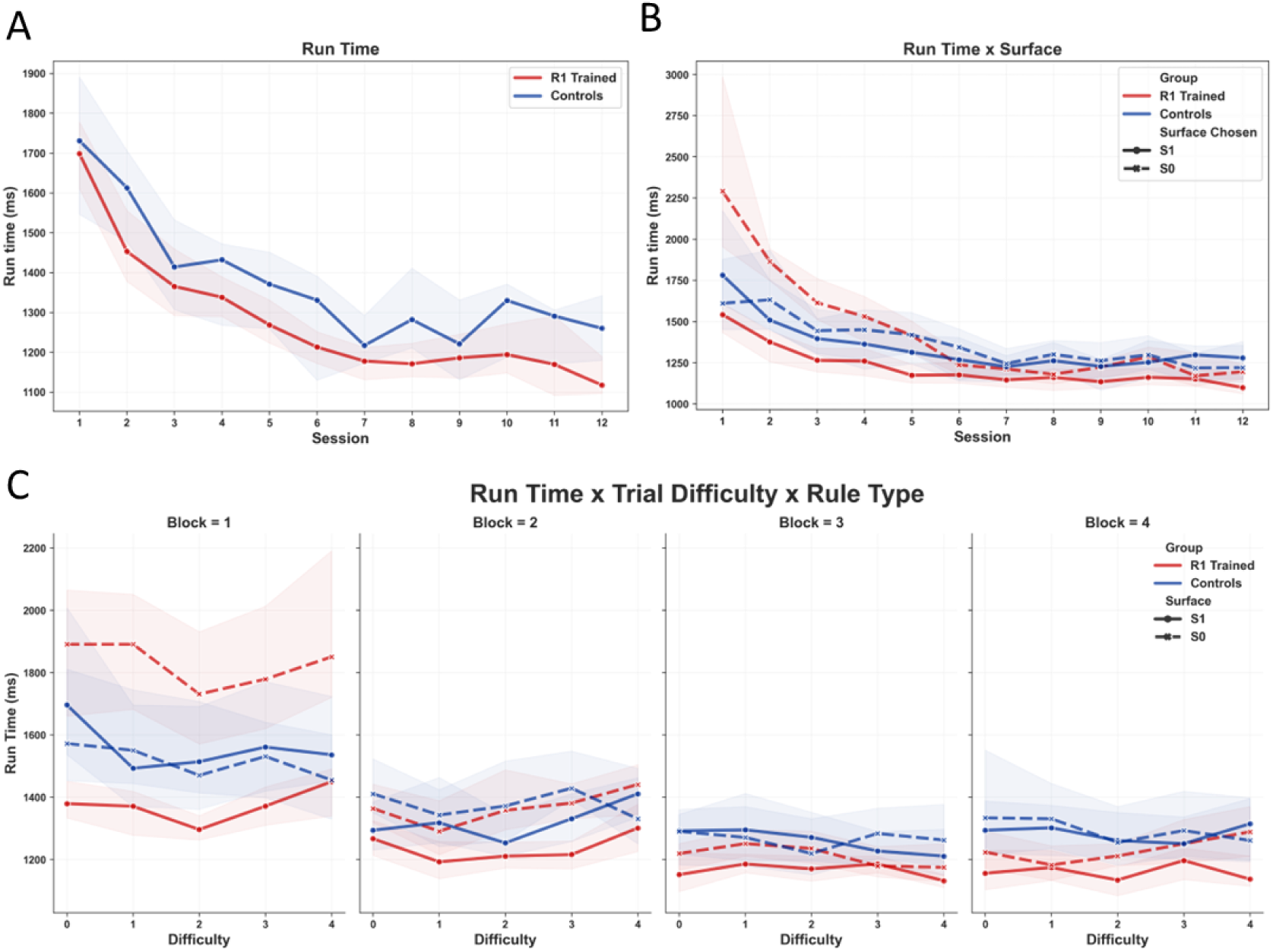
Confirmation bias-like effects highly persistent in post-decision response vigor/confidence. We furthermore developed analyses for measuring post-definitive decision run time as a proxy measure for response vigor/confidence. This behavior too proved to be highly revelatory with respect to R1 interference during outcome-anticipatory processes during EdRR. All error bands represent 95% confidence intervals (see main text for detailed statistical analyses). **(A)** Median run times (in milliseconds), independent of surface, by session. Averaged across all EdRR sessions, R1 trained animals displayed lower post-choice run times, an indicator of post-choice confidence, than controls. This was in contrast to the R1 training phase during which controls displayed lower overall post-choice run times. **(B)** Median run times (in milliseconds) by surface chosen. R1 trained animals displayed significantly higher per session S0 run times up until EdRR 5 (note the lag in group homogenization compared with evolution of decision latency). Furthermore, R1 trained run times on S1 arms remained consistently lower, indicating a certain level of persistent “over-confidence” in these choices, up until session 12. **(C)** Median run times (in milliseconds) by difficulty by block. Confirming the robustness of these lower S1 run times, we can see here that it was reliably the case over all trial difficulty levels and in all blocks.

To sum up these three sets of deliberative behavioral results, R1 interference during EdRR was evidently no simple affair of unreflective persistence of previous learning. Rather, when competitive relative to a present choice decision (notably S0-leaning choices), R1 interference appears to have *amplified* deliberation and it was in and through these very processes of boosted deliberation that the R1 bias most clearly manifested.

## Discussion

To our knowledge, the present study is the first to provide robust evidence of non-human animals manifesting myside confirmation bias-like behaviors during processes of cognitive deliberation. Such behaviors are called out by situations in which previously reinforced beliefs, understood as rules of action (Peirce, 1878) or state-action policies (Sutton & Barto, 2014), must be updated or revised in light of novel information (which can be either confirmatory or disconfirmatory with respect to priors) drawn from a complex and dynamic environment. Our claim is that the observations and analyses laid out in the present study constitute validation of a mouse model of just such real-world, everyday-like human situations of rule revision and cognitive resistance to the same. The introduction of this model therefore has two principal interests: 1) Its inherent interest as a means for gaining deeper understanding of how the mammalian mind-brain cognitively and neurophysiologically adapts to bias-generating situations: 2) Its pre-clinical research interest with respect to domains where differential confirmation bias-like phenotypes have been observed in humans, such as addiction rehabilitation and behavior change (Granero et al., 2020; Prochaska, 2008), schizophrenia (Doll et al., 2014), ageing (Wilson et al., 2018), as well as other physiological (Rollwage et al., 2020; Rollwage & Fleming, 2021; Stanovich et al., 2013; Stanovich & West, 2007) and pathological (Balzan et al., 2013) conditions. Additionally, this model validation in and of itself furthermore alters the research horizon with respect to our understanding of the neurobiological and environmental evolution and development of the higher reasoning faculties in humans.

### Biased deliberation: a representational interpretation

In keeping with our recent analyses of cognitive deliberation during active forgetting-dependent everyday-like memory (EdM) in mice (Stevens et al., 2023), our interpretation of the myside confirmation bias-like behaviors reported in the present study also posits subjectively experienced representation of cognitive content (without however making any claims as to the neural or phenomenological nature of the representations in question). As applied to the EdM protocol, the representational interpretation can be summed up as follows: 1) Deliberation is deliberation over representation (Redish, 2016): 2) Accurate anticipatory EdM responding relies on comparison and selective inhibition of competing spatiotemporal context-relevant representations (e.g. mnemonic episodes; Anderson & Hulbert, 2021): 3) Mnemonic episodes become spontaneously temporally organized via spatial context-induced repetition of active forgetting, the impact of which is to progressively weaken targeted representations (what is referred to as retrieval-induced forgetting, or RIF; Bekinschtein et al., 2018; Stevens et al., 2023). As far as empirical confirmation of this interpretation goes, in EdM trials entailing higher levels of prior RIF/representational weakening we have reliably observed both lower performance *and* decreased deliberation (Stevens et al., 2023).

Turning to the present mouse model of everyday-like rule revision (EdRR), our observations here suggest that bias from prior learning manifests via cognitive processes such as pattern completion (Horner & Burgess, 2014). By this process, putatively striato-hippocampal signals excited via active perception of R1 elements in the EdRR environment (e.g. surface type) could holistically reactivate cognitive-affective representations of recent R1-compatible memory episodes. Such holistic reactivation, we advance, could explain the *increased* deliberation observed in R1 trained animals during EdRR. Indeed, independently of the level of prior RIF entailed by a given trial, stably high levels of deliberation were observed both when animals *revised towards* an S1 choice (which often resulted in error) and while, conversely, they *executed* an S0 choice. We posit that the representational reactivations in question here could be either of episodes where the animal was *not* rewarded on the S0 surface, or of previous episodes where it had *revised* an initial S0 choice and was then rewarded on the S1 arm. By our interpretation, this R1-weighted representational reactivation during S0 choice execution would give rise to the deliberative hesitation and/or choice regret/revision which we saw. Conversely, deliberation either when animals revised towards an S0 or while they executed an S1 choice decreased normally as a function of increasing RIF (“normally” = replicating the deliberation profile observed both in controls as well as, previously, in naïve animals performing the EdM task (Stevens et al., 2023)). Moreover, under certain conditions R1 bias was also ‘positively’ visible in a heuristic-like manner (Kahneman, 2011), namely in the observation that definitive S1 choices were reliably and persistently executed faster/less hesitantly than definitive S0 choices. We therefore interpret the biased decision making observed as resulting from reactivation of representational content which is *excessively* weighted by prior associative cognitive-affective content, embodying strong figurative ‘No!’ or ‘Yes!’ signals with respect to the phases of an animal’s ongoing choice behavior on a given trial. Indeed, as RIF increases as a function of trial difficulty, this excessive weighting of R1-biased content becomes all the more disproportionate by comparison to a weakening episodic memory representational baseline.

These interpretations have implications both for the validity of the present model and for what our findings potentially reveal about the evolution of the neurocognitive processes underpinning myside confirmation bias behavior in humans. In decision making situations of ambiguous representational complexity (e.g. EdRR), perception of elements that resonate with strong prior beliefs/rules of action robustly gives rise to an increase in deliberative processes (where these processes represent what in humans we might call *reasoning*). However, since the deliberation involved is substantially fueled by *prior* cognitive-affective content internalized by the agent (i.e. idiocentric deliberation), as opposed to by accurate context-sensitive perception of the present environmental situation itself (i.e. allocentric deliberation), we advance that the biased behavior we observe in mice more closely parallels what in humans is referred to as *motivated reasoning* or *rationalizing*, a behavior that certain theories of the evolution of human cognition preclude in non-human species (Mercier & Sperber, 2017). Indeed, reflecting the lack of correlation between general intelligence and myside bias susceptibility documented in humans (Stanovich et al., 2013; Stanovich & West, 2007), simply ‘thinking more’ (i.e. deliberating) offers mice performing the EdRR task no observable protection from bias either. On this basis, we would suggest that a more essential difference between mice and men in pre- and post-choice cognition more likely consists in the powerful human capacity for outcome *reinterpretation*. This point will be more fully developed elsewhere (Stevens, forthcoming) but, in short, the cognitive mechanism referred to informally as ‘sour grapes’ (after the fable of the fox and the grapes found in both Aesop and La Fontaine) allows for absence of reward to be reframed and re-evaluated as a *positive* outcome, retroactively ‘rationalizing’ an immediately negative outcome into the most globally reinforcing cognitive option available, thereby protecting the integrity of prior beliefs and evading the need to revise them (Kaplan et al., 2016). Hence, although the ‘sour grapes’ approach to resolving cognitive dissonance (Festinger, 1957) may certainly *contribute* to how myside bias manifests specifically in humans, we would advance that it is better understood as an auxiliary to and not a core element of biased cognition.

### Impact on everyday-like memory performance from a previously acquired, partially antagonistic cognitive rule

When animals trained to reliably respond to the tactile stimulus-response rule (R1) were transferred to the EdRR environment, we observed that prior learning generated only a slight although initially significant negative impact on overall EdM performance compared to control animals (recall control condition = rewarded on every trial during R1 training regardless of surface chosen). Nevertheless, EdM performance in the EdRR environment was lower, even in controls, than performances typically observed in the classical EdM task with naïve animals (Marighetto et al., 2011; Stevens et al., 2023). It is important that this point be underlined: overall EdM performance *does* appear to be negatively affected by R1 training *and also* by our control condition. This points to a cognitive consequence of the control condition that is worth underlining. In the EdRR phase, both R1 trained *and* control animals were dealing with comparably radical, albeit distinct, changes to the response-outcome (R-O) reward contingencies they had become habituated to in their respective training phase environments. This therefore functioned as a control for any initial frustration the R1 population was likely to experience in the EdRR environment upon not obtaining reward when expected (Amsel, 1958; Martín-García et al., 2015), an interesting phenomenon in itself but not the direct object of our investigations. However, based on the fact that R1 trained animals were choosing S1 at an average rate of 83% during the final R1 training sessions, one might have expected them to initially continue to choose S1 arms at a comparable rate in the EdRR environment and, therefore, have expected their initial EdM performance to fall significantly *below* chance level. Why did this not happen?

### EdM performance and the impact of novelty on exploration

The links between novelty and exploration in both humans and rodents are well established and have been the subject of much research and discussion (Farahbakhsh & Siciliano, 2021; Lustberg et al., 2020; McDonald et al., 2004; Park et al., 2021). In our experiments, upon initial introduction to the EdRR environment, which we had expressly made as sensorially novel as possible with respect to the R1 environment (new room, new radial maze, new lighting conditions, new ambient odor), not only did the R1 trained population not immediately prioritize exploration of S1 arms, they pointedly did the opposite, a significant majority preferring to explore S0 arms on the first two trials of the first session of EdRR. Recalling furthermore that during EdRR only the S1 arm was rewarded on the initial presentation of each of the three pairs, a majority of animals who chose S0 on the first EdRR trial (and were therefore *not* rewarded, thus constituting a *confirmation* of R1) nevertheless chose S0 again on the second trial. This result was as surprising as it is fascinating since it indicates that environmental novelty can drive not only topographical exploration of the new environment but also *nomological* exploration, i.e. exploration directed precisely beyond the bounds defined by an internalized behavioral rule. It is moreover possible that these two modes of exploration are connected, potentially subsumed under a global information gain exploratory drive (Inglis et al., 2001). This is a question we intend to explore in future work as part of a computational modeling approach aimed at deeper understanding of the behaviors observed in this study. For control animals, absence of reward on an initial S0 choice during EdRR constituted the first time they had encountered a no-reward trial outcome of any kind. Thus, beyond the environmental novelty, it is possible to posit a further effect of frustration or surprise at play (Amsel, 1958; Xu et al., 2021). This could explain why, of those controls who chose S1 on the first trial of EdRR 1, only half (i.e. commensurate with random choice behavior) chose S0 on the second trial, whereas of those who chose the *un*rewarded S0 on the first trial, a *majority* chose S0 again on the second trial, revealing a higher probability for an initial and counter-intuitive ‘lose-stay’ strategy which we do not believe can be satisfactorily explained by decision inertia (Alós-Ferrer et al., 2016) and therefore merits further investigation.

Manifestation of the novelty-induced exploratory drive was not however limited to just the first two trials of EdRR 1. The median number of trials taken by the R1 population to at least once explore all six available arms, S1 and S0, of the EdRR radial maze approached optimal level (i.e. 7 trials) during EdRR 1, making them stronger initial explorers even than controls. Since we hypothesized that this burst of exploration was driven primarily by the salient novelty of the EdRR environment, we also predicted that we would see less striking exploratory behavior later in EdRR 1 and less again in EdRR 2. This prediction was borne out by the data. First, on all trials (*post* the initial three) of EdRR 1, the R1 population was more likely to choose S1, with the surface choice probability shifting back towards S0 only on the final trial which, tellingly, was a difficulty level 0 trial. Subsequently, we observed that R1 trained mice took significantly more trials to explore all available arms at least once during EdRR 2 compared to EdRR 1, fitting with our hypothesis that the initial exploratory boost was primarily a cognitive response to novelty. As mentioned above, there is some precedent in the literature demonstrating a facilitation of S-R reversal learning when this takes place in a novel context rather than in the same context as the initial learning (McDonald et al., 2004). However, our EdRR protocol was designed precisely *not* to be a simple reversal paradigm, and while previous results in rule reversal tend to follow a steady upwards curve, here we notably saw no mean difference in P(S1) in experimental animals between EdRR 1 and EdRR 2, despite an average of ∼60% of all EdRR 1 trials giving rise to R1 *dis*confirmations. Thus novelty itself, and not repeated errors from behaving according to R1, was the primary explanatory factor behind initial exploratory, i.e. S0 directed, choice behavior. What these results seem to reveal is that the R1 population had no deficit with respect to initial exploratory ‘sampling’ of a novel environment but *were* subsequently impaired in evaluating these novel context experiences and using them to appropriately update their level of adherence to the prior R1 rule. Finally, if we compare this to other recent work which demonstrated, in naïve mice, that increased novelty improves consolidation of proximal experiences (Takeuchi et al., 2016), we get an even better idea of just how strong the impact of myside confirmation bias-like effects from prior learning can be, since the manifestation of the bias we observed seems precisely to relate to a failure to adequately consolidate R1 disconfirmation experiences between one session and the next (24 hours later).

### Within-error R1 bias

While only a slight and initial significant difference was observed between the R1 and control populations in overall numbers of session-by-session EdM errors, the type or quality of the errors committed by the two groups was vastly different. On average, the R1 population committed almost 90% of their errors in EdRR 1 on S1 arms compared to an almost even S1/S0 split in errors observed in controls. Curiously however, we found no correlations between R1 training phase performances (whether final, summed, or weighted) and R1 error bias, nor even between EdRR 1 exploration score and R1 error bias. This was all further indication to us of the multiplicity of cognitive processes mobilized by the EdRR protocol, notably those that inhibit the innate exploratory drive during R1 training plus those involved in sensitivity to environmental novelty and subsequent inhibition of interfering/intrusive prior cognitive content.

However, by block 4 of EdRR training (sessions 10-12), the within-error R1 bias index no longer differed between the R1 trained and control populations, with both groups displaying a slight within-error R1 bias. Naturally this raised the question of why the controls had developed such a bias during EdRR training. One potential explanation is the contingent fact that, for the duration of EdRR, the session-by-session probability of control animals receiving a reward having chosen an S1 arm, P(Rw|S1), was reliably higher than the probability of receiving a reward having chosen an S0 arm, P(Rw|S0). Thus some form of putatively Bayesian striatal process working in the background to statistically track and encode R-O values from session to session could explain a subsequent emergent trend for the more often rewarded surface to be preferred (Kim et al., 2009; Samejima et al., 2005) especially on more difficult EdM trials. By contrast, in the case of the R1 population, P(Rw|S1) was significantly *lower* than P(Rw|S0) until block 4 of EdRR, so that their within-error R1 bias persisted not because of but *despite* their experiences in the EdRR environment.

### Conclusion: towards a multiple memory systems conception of myside confirmation bias

Reflecting classic studies on multiple memory systems (McDonald & Hong, 2004; McDonald & White, 1993; White & McDonald, 2002), the results validating the EdRR model of myside confirmation bias suggest that the EdM and especially EdRR protocols are uniquely capable of differentially mobilizing up to four interacting (competing and/or cooperating) memory systems: episodic memory (EM), striatal memory, affective memory, and working memory (WM, including inhibitory control). As already mentioned, it is widely accepted that Y- and T-maze spatial alternation preferentially engages WM (Albayram et al., 2016; Aultman & Moghaddam, 2001; Jobson et al., 2021; Shoji et al., 2012). In the classical EdM task also, animals with either lesion of the CA1 region of the hippocampus (Marighetto et al., 2011) or genetically induced EM dysfunction (Stevens et al., 2023) are capable of above chance level performance on level 0 trials only. These results suggest that hippocampal EM is *necessary* for level 1 to level 4 trials but not for level 0 trials, a conclusion that corroborates well-established hypotheses which take WM to be a cortical rather than hippocampus-centered memory system (Baddeley, 2003; Curtis & D’Esposito, 2003; Lara & Wallis, 2015). Conversely, the performances of transgenic and aged animals (Stevens et al., 2023) also demonstrate that the hippocampus is not *sufficient* for success on level 1 to 4 EdM trials; top-down cortico-hippocampal active inhibition of competing cognitive content is also required.

During high difficulty EdRR trials, we observed that the R1 population was significantly more likely to respond according to the (putatively) striatally acquired stimulus-response R1 rule than according to the spontaneous exploratory spatial alternation EdM rule. It is therefore plausible to suggest the following mnemonic model of the behavior: as *contextualized* cortico-hippocampal representational memory systems lose representational strength, due to repeated RIF, action choice will increasingly fall to *decontextualized* S-R striatal and/or affective amygdalar memory systems. If striatal R-O values are indeed updated in some kind of ongoing Bayesian manner (Ballard et al., 2018; Kim et al., 2009; Markowitz et al., 2018; Nonomura et al., 2018; Samejima et al., 2005), incorporating the full, context *in*dependent historic of P(Rw|S1) from both the EdRR *and* prior R1 environments, and if the striatum is more heavily relied on during action selection on more difficult trials, then this could explain why P(S1) remains significantly higher on high difficulty trials compared to easier ones even into block 4 of EdRR. Our control group data also bring some convergent evidence to this multiple memory systems hypothesis. In controls, P(Rw|S1) was reliably *higher* than P(Rw|S0) in all sessions. In a breakdown of both P(Sn) and within-error R1 bias as a function of trial difficulty, we then saw that controls developed a significant trend towards higher P(S1) on level 4 trials and also committed a significantly higher proportion of S1 errors on level 4 trials in blocks 2 to 4. Hence, taken together, both sets of data suggest that resorting to a striatal, statistical response strategy on trials entailing high RIF is a spontaneous cognitive strategy. We did not, however, observe any sign of a putative affective/amygdalar contribution in either the run time or choice revision behaviors of control mice, potentially indicating that affective circuits are less sensitive to subtle statistical differences in R-O values. This is a question for further investigation. If such an ongoing surface-based statistical evaluation is indeed taking place during the EdRR phase, then rule revision with respect to R1 can be seen as a case of taking one step backward for every two steps towards overcoming R1 interference. This would seriously delay the updating process compared to, for example, more classical all-or-nothing reversal learning protocols and could therefore be a key component in the delay we observed in R1 trained animals with respect to their updating of P(Sn) as a function of both P(Rw|Sn) and the balance of R1 confirmations versus disconfirmations (where per session R1 disconfirmations significantly outweighed R1 confirmations during the first 5 sessions of EdRR).

Finally, striatal memory would not play a role *only* in the most difficult trials. Since the striatum is essential to the selection and initiation of movement, it is ultimately involved in decision-making, and especially decision *execution*, at all levels of difficulty. Indeed, in the R1 population, striatal S-R responding leads to significant interference at all trial difficulty levels. The role of the striatum in action choice has previously been modelled as a softmax action selection rule (Kim et al., 2009) whereby the probability of choosing one action over another (e.g. ‘explore arm on the left’ vs ‘explore arm on the right’, or ‘respond to S1’ vs ‘respond to S0’) varies according to the difference between the values attributed to each of the potential actions. This can be expressed as *a* = (*Q*_S1_ – *Q*_S0_), where *a* stands for the chosen action and *Q* stands for the estimated/predicted value of each action option (e.g. *Q*_S1_ = estimated/predicted value of choosing S1), as per the *Q*-Learning reinforcement learning approach (Watkins & Dayan, 1992). In this kind of picture, what we suggest is that on easier difficulty trials, signals coming from cortical working memory and/or hippocampal episodic memory to the striatum are low in uncertainty/noise. These low uncertainty signals would have the capacity to tip the *Q*_S1_ vs *Q*_S0_ balance away from the striatum’s local and historical valuations of these options. However, as RIF-induced uncertainty increases, the signals coming from these upstream memory systems would become weaker and noisier and thus less able to influence striatal action selection. The same can be suggested with respect to a putative amygdalar contribution to deliberative decision making, especially during the post-initial choice phase wherein the relevant parameters (i.e. choice revision and run time) tend to be most pronounced on easier trials in classical EdM but not in EdRR, where they are equally pronounced across all trial difficulty levels.

If, as we suggest, such a multiple memory systems interpretation of the confirmation bias-like behavior we have experimentally observed is accurate, then the basis of what psychologists refer to as myside bias may fundamentally be a latent neurocognitive fact of all organisms having evolved analogous multiple memory systems. Through this interpretative lens, the particular competitive combinatory activation of these systems (which in humans we call myside confirmation bias) becomes something that has been called out or co-opted (Gould et al., 1979) into existence by the specific epistemic environments of belief, knowledge, persuasion, and often dogmatic education which, over millennia, we have created for ourselves to inhabit. We believe the present study opens the way to much future research in this direction across a spectrum from comparative neurophysiology to developmental and educational psychology.

## Supplementary figures

**Supplementary figure S. 1.**
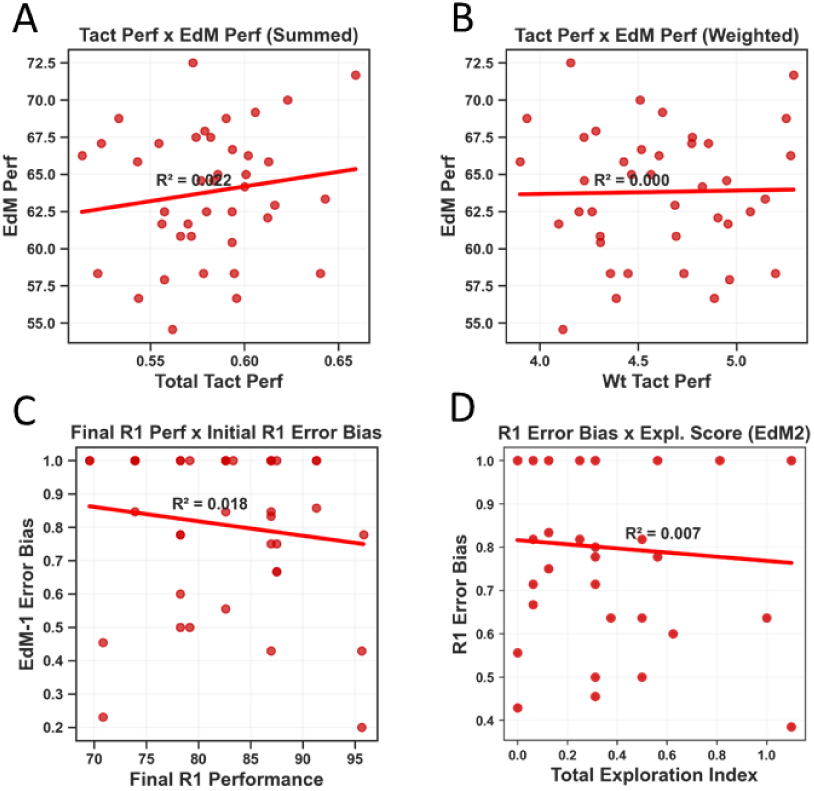
Absence of correlations hints towards interaction of multiple cognitive systems. Several linear regression analyses were calculated from individual R1 trained population parameters in order to detect correlations between behaviors which may intuitively have been correlated. **(A)** No correlation was found between summed tactile discrimination (R1) performance in the training phase and summed EdM performance during EdRR (a negative correlation might have been expected). **(B)** The R1 performance was then weighted, discounting earlier compared to later performances, on the eventuality that earlier low R1 performances were occluding a relationship. No correlation between these values was observed. **(C)** Specifically between R1 performance in the final R1 session only and within-error R1 bias during the first EdRR session only (two sessions separated by 24 hours), we again found no correlation. Notably, both relatively weak and strong R1 performers were just as likely to display a 100% S1 error bias during EdRR 1. **(D)** Finally, having observed that novelty per se had a major impact on behavior, especially during EdRR 1, we instead compared individual EI values in EdRR 2 with levels of within-error R1 bias also in EdRR 2. No correlation was observed.

**Supplementary figure S. 2.**
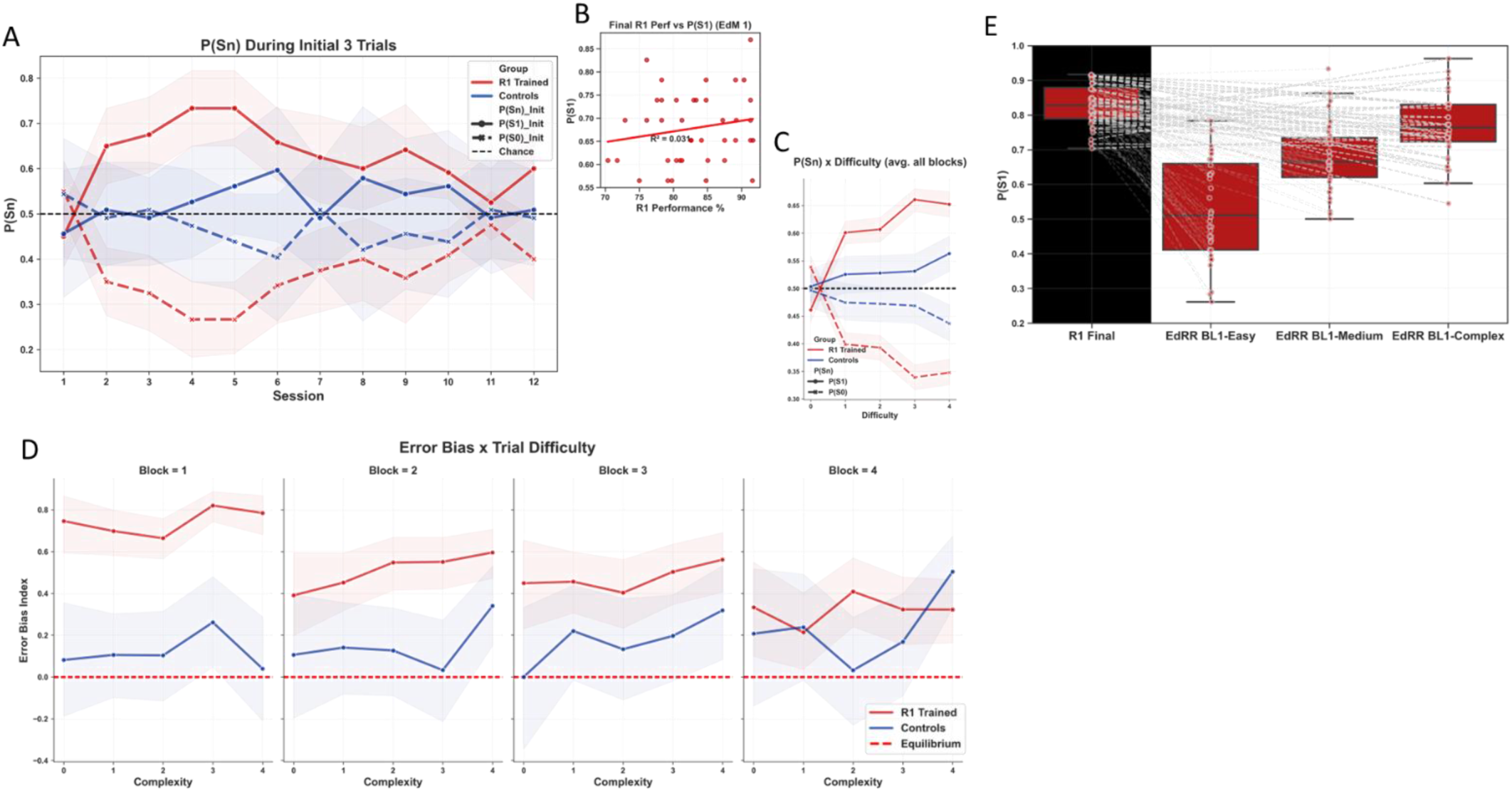
S1 bias clear and robust during EdRR, as a relative difference within errors and as a function of trial complexity in overall P(S1). All error bands represent 95% confidence intervals, vertical spaces between bands indicate statistical significance (detailed statistical analyses in main text). **(A)** Isolated P(Sn) values for initial 3 trials not included in analyses according to trial complexity. The impact of novelty on initial exploratory behavior in R1 trained animals during EdRR 1 is clearly observable. Following EdRR 1, P(S1) is significantly higher than P(S0) in almost all sessions in the R1 trained population. In controls, we observed more fluctuation, but still with a slight tendency for P(S1) to be higher. Recall that in these initial 3 trials, i.e. the first trial on each pair, only S1 was rewarded, thus P(Rw|S1) = 1 and P(Rw|S0) = 0 for these trials. **(B)** Linear regression analysis found no correlation between final R1 performance and overall P(S1) in EdRR 1. **(C)** P(Sn) as a function of trial complexity averaged across all sessions/blocks. In this representation, it is clear that P(S1) is, on average, slightly higher in controls on all trial complexities above level 0, but this reaches significance only on level 4 trials. **(D)** As we had done for the overall within-error R1 bias index, we also calculated the relative difference in S1 versus S0 errors as a function of trial complexity. At this level, there was no dependence of within-error R1 bias on trial complexity, only a slight dependence on time/repeated training which mirrored the trend of the overall within-error R1 bias to decrease over time. However, in controls, we can again see that, in blocks 2-4 especially, this population was significantly more likely to make errors on S1 than on S0 arms on complexity level 4 trials. **(E)** Having seen from figure 4 that P(Sn) in R1 trained animals seemed to mark out 3 distinct groupings of complexity (level 0, easy; levels 1-2, medium; levels 3-4, complex), we looked at P(S1) in the final R1 session (which was equal, by definition of the task, to R1 performance) and compared it to P(S1) values across the first block of 3 EdRR sessions grouped according to these three levels. This provides a clear visual representation of just how close R1 trained animals were in early EdRR sessions to performing on complex trials almost identically to how they had performed in the R1 environment where they had never experienced any R1 disconfirmations.

**Supplementary figure S. 3.**
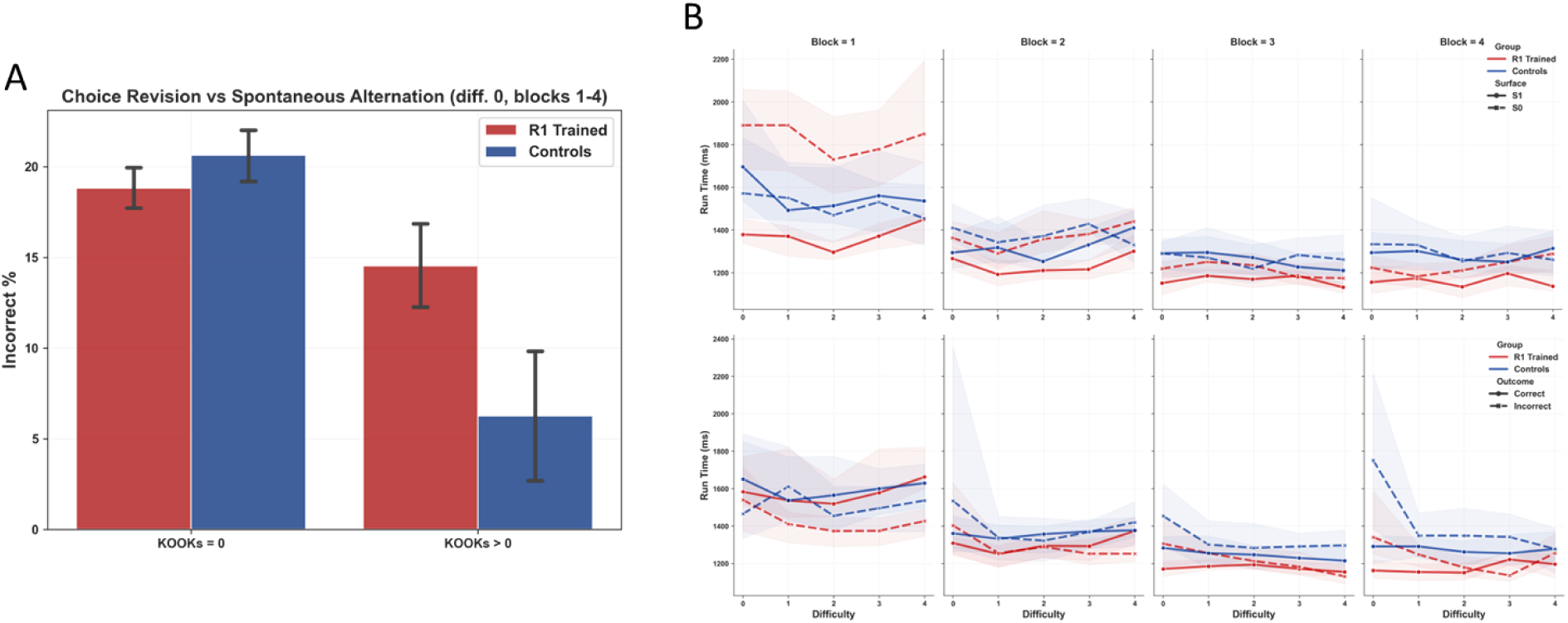
Interactions of surface, outcome, and deliberation. All error bands represent 95% confidence intervals. **(A)** In level 0 trials, almost all choice revision in controls was rectifying whereas R1 trained animals were comparatively likely to make an error with as without choice revision. 88% of these errors with choice revision, averaged across all blocks, were from S1 final choices (see main text). **(B)** Top; median run time by surface by difficulty by block. **(B)** Bottom; median run time by outcome by difficulty by block. As shown above, R1 trained animals maintain persistently lower run times on S1 in all blocks and at all trial complexity levels. As a function of outcome, we can see that controls in EdRR more quickly develop a stable run time phenotype across all trial complexities (i.e. higher run times on incorrect outcome trials; see supplementary figures A.3 + A.6) compared to R1 trained animals, which show a lag in this respect on more complex trials especially. This could, precisely, be caused by a sustained “over-confidence” effect on S1 choices, independently of outcome.

